# State-space optimal feedback control of optogenetically driven neural activity

**DOI:** 10.1101/2020.06.25.171785

**Authors:** M F Bolus, A A Willats, C J Rozell, G B Stanley

## Abstract

**Objective:** The rapid acceleration of tools for recording neuronal populations and targeted optogenetic manipulation has enabled real-time, feedback control of neuronal circuits in the brain. Continuously-graded control of measured neuronal activity poses a wide range of technical challenges, which we address through a combination of optogenetic stimulation and a state-space optimal control framework implemented in the thalamocortical circuit of the awake mouse.

**Approach:** Closed-loop optogenetic control of neurons was performed in real-time via stimulation of channelrhodopsin-2 expressed in the somatosensory thalamus of the head-fixed mouse. A state-space linear dynamical system model structure was used to approximate the light-to-spiking input-output relationship in both single-neuron as well as multi-neuron scenarios when recording from multielectrode arrays. These models were utilized to design state feedback controller gains by way of linear quadratic optimal control and were also used online for estimation of state feedback, where a parameter-adaptive Kalman filter provided robustness to model-mismatch

**Main results:** This model-based control scheme proved effective for feedback control of single-neuron firing rate in the thalamus of awake animals. Notably, the graded optical actuation utilized here did not synchronize simultaneously recorded neurons, but heterogeneity across the neuronal population resulted in a varied response to stimulation. Simulated multi-output feedback control provided better control of a heterogeneous population and demonstrated how the approach generalizes beyond single-neuron applications

**Significance:** To our knowledge, this work represents the first experimental application of state space model-based feedback control for optogenetic stimulation. In combination with linear quadratic optimal control, the approaches here should generalize to future problems involving the control of highly complex neural circuits. More generally, feedback control of neuronal circuits opens the door to adaptively interacting with the dynamics underlying sensory, motor, and cognitive signaling, enabling a deeper understanding of circuit function and ultimately the control of function in injury or disease.

## 1. Introduction

Over the last two decades, there has been a rapid expansion of tools and technologies for recording the large-scale activity within and across brain structures at single neuron resolution ([1, 2]). In parallel, the development of optogenetics provided the ability to optically excite or inhibit neural activity in a cell-type specific manner ([3]). Together, these advances in the ability to ‘read’ or ‘write’ the neural code have led to a wide range of discoveries of the circuit mechanisms underlying sensory, motor, and cognitive processes ([4]). The integration of recording and optogenetic stimulation techniques, however, has received comparatively little attention until recently ([5–13]; for review [14]), and in most cases these closed-loop systems utilize event-triggered or on-off control rather than continuous feedback. While feedback control is the engineering cornerstone for the function of a wide range of complex technologies ranging from communication to flight, applying this perspective in the nervous system remains more theoretical ([15–19]) than experimental. In this study, we establish a general framework for continuously-modulated closed-loop optogenetic control of neuronal circuits, where optical actuation is determined in real-time by comparing measured neuronal spiking to target activity. This work opens up possibilities for investigation of poorly understood mechanisms of the underlying circuitry and for adaptively interacting with the circuit dynamics within and across brain regions that constantly change in response to the internal and external environments.

Electrical stimulation has been the gold standard for manipulating the activity of neurons at fast time-scales, and remains the basis for clinical interventions like deep brain stimulation ([20]). However, this approach suffers from lack of specificity while also typically precluding simultaneous measurement of the activity of the neurons being stimulated. While not yet clinically viable, optogenetics offers an alternative approach that enables cell-type specificity, bi-directional actuation, the ability to simultaneously stimulate and obtain electrophysiological recordings, and a potentially lesser degree of unnatural synchronization of the local population ([21]). This presents an attractive toolbox for the development of continuous, feedback control strategies where stimulation is continuously modulated based on real-time measurements of the local neuronal activity. There has been a range of studies where previously-determined stimulation is triggered based on recorded activity in a reactive closed-loop fashion ([8–11]). In addition to event-triggered control, a recent study has also used on-off closed-loop control to gate photostimulation when recorded neuronal activity was below target levels ([13]). Although these approaches to stimulation have proven effective for their uses, they are fundamentally different from the continuously-graded feedback control we describe here, where stimulation is updated on a moment-by-moment basis as a function of the current and past measured neural activity. In previous studies, we have developed and demonstrated strategies for closed-loop optogenetic control of spiking activity of neurons in a cultured network and single neurons *in vivo* in the anesthetized brain ([5, 7]). While laying the conceptual groundwork, these approaches do not scale well to neuronal populations and do not take advantage of more modern approaches in control theory. Additionally, these previous studies had not yet applied optogenetic control in the context of wakefulness.

In this study we bridge the gap between optogenetics and established paradigms of more modern control theory by utilizing state-space models to capture single- and multi-neuron responses to optogenetic stimulation and employing optimal control to design the control loop for driving desired neuronal activity. Specifically, precise manipulation of neurons was carried out via optical activation of the excitatory opsin channelrhodopsin-2 (ChR2) expressed in the somatosensory thalamus of the awake, head-fixed mouse.

A feedback controller updated light intensity in real-time based on simultaneous electrophysiological measurements of the thalamic neurons being manipulated. A linear dynamical system model structure was used to approximate the light-to-spiking input-output relationship in both single-neuron as well as multi-neuron scenarios in cases where multiple neurons were measured simultaneously using multielectrode arrays. These linear state-space models were used in combination with linear quadratic optimal control to design feedback controller gains for the purpose of regulating thalamic firing around a desired target rate. The models were also used online for estimation of state feedback, using a parameter-adaptive Kalman filter for robustness to model-mismatch. The resulting controller-estimator feedback loop was deployed experimentally by way of a custom-written program running in real-time. This control scheme provided effective optogenetic control of firing rate in the awake brain, owing to the robustness to model accuracy granted by a parameter-adaptive Kalman filter that estimated a stochastically-varying process disturbance. Feedback control using this estimator resulted in very good firing rate tracking experimentally for the single neurons whose activity was used for feedback. By comparison, control was not as effective for other simultaneously-measured neurons not used for feedback. To investigate the generalizability and efficacy of these methods for future multi-output control scenarios, we demonstrate their application to multi-neuron feedback control of a population in simulation.

## 2. Methods

### 2.1. Animal preparation

All procedures were approved by the Institutional Animal Care and Use Committee at the Georgia Institute of Technology and were in agreement with guidelines established by the NIH. Experiments were carried out using either C57BL/6J mice that were virally transfected to express channelrhodopsin-2 (ChR2) or by single-generation crosses of an Ai32 mouse (Jax) with an NR133 cre-recombinase driver line (Jax) which grants better specificity of ChR2 expression in ventral posteromedial/posterolateral thalamus ([22]). In the case of viral transfection, ChR2 expression was targeted to excitatory neurons in the thalamus via stereotactic injection relative to bregma (approximately 2 × 2 × 3.25 mm caudal × lateral × depth) using 0.5 μL of virus (rAAV5/CamKIIa-hChR2(H134R)-mCherry-WPRE-pA; UNC Vector Core, Chapel Hill, NC) at a rate of 1 nL/s.

At least three weeks prior to recordings, a custom-made metal plate was affixed to the skull for head fixation and a recording chamber was made using dental cement while the animals were maintained under 1–2% isoflurane anesthesia ([23]). After allowing a week for recovery, mice were gradually habituated to head fixation over the course of at least five days before proceeding to electrophysiological recordings and optical stimulation. On the day of the first recording attempt, animals were again anesthetized under 1-2% isoflurane and a small craniotomy (1-2 mm in diameter) was centered at approximately 2 × 2 mm caudal and lateral of bregma. The animals were allowed to recover for a minimum of three hours before awake recording. At the time of recording, animals were head-fixed and either a single electrode or an electrode array coupled to an optic fiber (Section 2.2) was advanced through this craniotomy to a depth between 3-4 mm for thalamic recording and stimulation. Between repeated recording attempts, this craniotomy was covered using a biocompatible silicone sealant (Kwik-Cast, WPI). Following termination of recordings, animals were deeply anesthetized (4–5% isoflurane) and sacrificed using a euthanasia cocktail.

### 2.2. Experimental setup

All optical stimuli were presented deep in the brain via a 200 or 100 μm diameter optic fiber attached to a single tungsten electrode (FHC) or to a 32-channel NeuroNexus optoelectric probe in a 25 μm-spaced ‘poly3’ configuration (A1×32-Poly3-5mm-25s-177-OA32LP, NeuroNexus Technologies, Inc.), respectively. Command voltages were generated by a data acquisition device (National Instruments Corporation) in a dedicated computer running a custom-written RealTime eXperimental Interface (RTXI, [24]) program at 1 ms resolution. Command voltages were sent to a Thorlabs LED driver, which drove a Thorlabs M470F3 LED (470 nm wavelength blue light) connected to the 100-200 μm optical fiber. A commercially available data acquisition device and processor (Tucker Davis Technologies RZ2) measured extracellular electrophysiology. This system was used for single-channel PCA spike sorting, binning, and sending these binned spike counts at 2 ms resolution to the computer running RTXI over ethernet via UDP. The computer running RTXI for realtime control listened for datagrams over ethernet and linearly interpolated from 2 ms to the operating resolution of 1 ms. All told, the closed-loop processing loop was approximately 10 ms.

As mentioned above, the control and estimation algorithms were carried out in real-time at 1 ms resolution using a custom-written program. The program consisted of an RTXI ‘plugin’ linked against a C++ dynamic library that was responsible for online estimation of state feedback (Section 2.5) and the generation of control signals (Section 2.6). This functionality was provided as part of a state-space controller C++ class. The RTXI plugin forwarded the reference, or target, firing rate, model parameters, and feedback controller gains to a state-space controller object, and the controller returned an updated control signal each time it was queried by RTXI. This control signal was then routed by RTXI to the LED driver via a DAC (see above). All linear algebra was carried out using the C++ library Armadillo ([25]).

### 2.3. Offline spike sorting

For online control applications, single-channel PCA-based spike sorting was carried out in real-time using a commercially available electrophysiology system (Tucker Davis Technologies RZ2). Beyond tetrode recordings, spike sorting from high-density electrode arrays requires a multi-step process that is not feasible within the timescale of experiments with head-fixed awake animals. Kilosort2 ([26]) was used for all offline spike sorting, including single-channel recordings, in which case spatial whitening and common mode referencing steps were disabled. Initial sorting by Kilosort2 was then manually curated (additional merging/splitting of clusters) using the phy viewer. After manual curation, any clusters that met the following criteria were considered single units and used in this study: sub 1 ms ISI violations of *<*0.5%, sub 2 ms ISI violations of *<*2%, and mean waveform amplitude-to-standard-deviation ratio *>*4.

### 2.4. Mathematical modeling

Linear and Gaussian state space models were used in designing feedback controller gains before experiments as well as during the experiment as part of the state feedback estimator. These models were fit offline to neuronal data before experimental application of optogenetic control and were fit at 1 ms time resolution, which was also the operating resolution of the RTXI software used for real-time control and estimation during experiments. In addition to the single-unit quality selection criteria in Section 2.3, models were only fit to putative single units (called ‘neurons’ hereafter) whose activity was significantly modulated by optical stimulation. Following Sahani and Linden ([27]), a neuron’s response was considered significantly modulated if the amount of ‘signal power’ in the response was greater than one standard error above zero. Note that Sahani and Linden ([27]) define ‘power’ as the variance in time. We will refer to ‘signal power’ as ‘signal variance’ in this study.

The underlying dynamics of neural activity were approximated as a linear dynamical system (LDS) in which a number of latent ‘state’ variables, represented as the vector **x** ∈ ℝ^*n*^, evolve linearly in time:

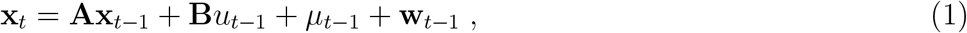

where *u*_*t*_ ∈ ℝ^1^ is the optical stimulus at time *t, μ*_*t*_ ∈ ℝ^*n*^ is a process disturbance, **w**_*t*_ ∼ 𝒩 (0, **Q**) is Gaussian noise of covariance **Q, A** ∈ ℝ^*n*×*n*^ is the state transition matrix, and **B** ∈ ℝ^*n*×1^ is the input vector (generally a matrix). Note that the disturbance, *μ*, was assumed to be zero during model fitting. However, for robustness in control applications, *μ* was allowed to be non-zero and to vary stochastically over time for the purpose of online state estimation (Section 2.5.2).

#### 2.4.1. Gaussian linear dynamical system

Linear and Gaussian models were used for control system design and implementation because of the relative simplicity and ubiquity of linear control approaches. In this case, the output of an LDS **y** ∈ ℝ^*p*^ is modeled as a linear transformation of a latent state **x** and is assumed to be corrupted by additive Gaussian noise before measurement in the form of binned spiking, **z** ∈ ℝ^*p*^:

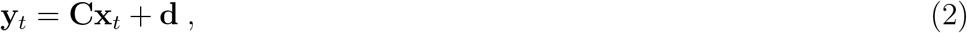

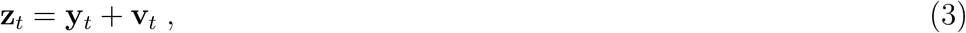

where **C** ∈ ℝ^*p*×*n*^ is the output matrix, **d** ∈ ℝ^*p*^ is an output bias term that describes the baseline firing rates of the *p* outputs (here, neurons), and **v**_*t*_ ∼ 𝒩 (**0, R**) is zero-mean Gaussian measurement noise of covariance **R** ∈ ℝ^*p*×*p*^. As a system whose dynamics evolve linearly and whose observation statistics are assumed to be additive/Gaussian, this is termed a Gaussian LDS, or GLDS ([28]). The bias term **d** was estimated as the average firing rate of each channel during spontaneous periods without optical stimulation, and GLDS models were fit relating *u*_*t*_ and (**z**_*t*_–**d**) using subspace identification (N4SID algorithm, [29]).

#### 2.4.2. Poisson linear dynamical system

While GLDS models were used for control and estimation, we evaluated their performance in capturing light-driven firing rate relative to a spiking model. As it is a more accurate statistical observation model for spike count data, we fit linear dynamical systems with Poisson observations, so-called Poisson LDS, or PLDS ([28, 30]). In this case, the underlying latent state(s) of the LDS is mapped to an output firing rate by a rectifying exponential nonlinearity and the measured spike counts are assumed to be drawn from a Poisson process driven at the given rate:

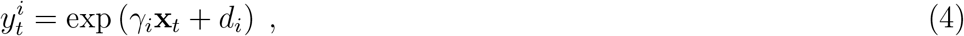

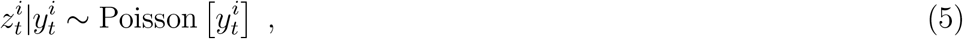

where 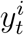 is the firing rate and 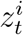 the measured spike counts of the *i*^th^ output at time *t*. For the purposes of this study, PLDS models were fit by first estimating a GLDS model. The row vectors *γ*_*i*_ that describe the log-linear contributions of each state to output firing rates were assumed to be scaled versions of the GLDS output matrix rows: *i.e*., for the *i*^th^ output,

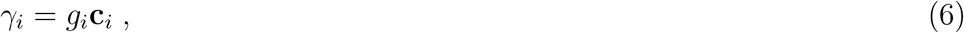

where **c**_*i*_ is the corresponding row of the GLDS output matrix **C**.

Note that at each time point the outputs are statistically independent conditioned on the state, allowing the output function parameters to be estimated in an output-by-output fashion. The resulting 2*p*-parameters of the PLDS output function were fit by maximizing the log-likelihood of the model one output at a time, given the predicted state sequence:

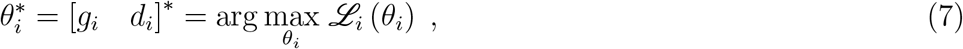

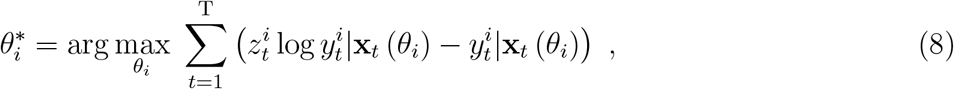

where 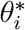 are the parameters and ℒ_*i*_ the log-likelihood of the model for the *i*^th^ output, (·)^∗^ denotes the result of the optimization, and

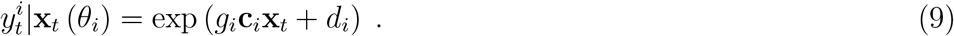

This optimization was carried out iteratively until parameter convergence for each output by analytically solving for *d*_*i*_ and numerically solving for *g*_*i*_ using Newton’s method in a manner analogous to Smith *et al* ([30]).

#### 2.4.3. Finite impulse response model

While state-space models were used in this study, finite impulse response (FIR) models were also fit in order to provide empirical estimates of the light-to-spiking responses that did not depend on choices such as number of latent states. Moreover, FIR models, often termed (‘whitened’) spike-triggered average (STA) models, are widely used to characterize neuronal responses to stimuli ([31]), so they are are a more familiar model type for much of the neuroscience community and provide a useful point of comparison for state-space models which are less frequently used in this context. Contrary to state-space models whose outputs share a set of dynamical states, in FIR models the optical stimulus (*u*) is related to the output firing rates (**y**) of *p* neurons in the following manner:

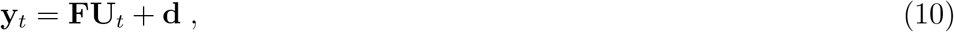

where **U**_*t*_ is a *q*-dimensional column vector of stimulus history up to time step *t* inclusive,

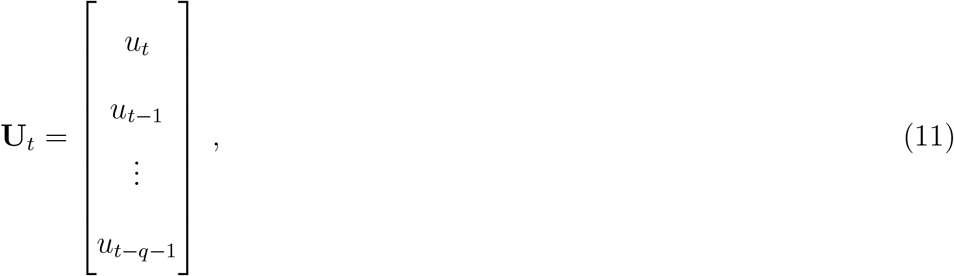

and **f**_*i*_ is the impulse response of the *i*^th^ output, comprising the rows of **F**, and **d** is the output bias as before in the case of the GLDS model. Note that this is effectively a convolution of a set of *p* FIR filters with the stimulus. The output **y** is assumed to be corrupted by additive Gaussian noise before being observed/measured in the form of binned spiking, **z**:

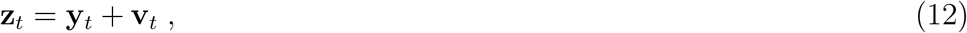

where **v**_*t*_ is the measurement noise as described in the case of the GLDS model previously. This FIR model was fit by ordinary least squares linear regression between (**z**_*t*_ − **d**) and corresponding 100 ms stimulus histories, *i.e*., **U**_*t*_ ∈ ℝ^100^ at Δ = 1 ms sample period.

#### 2.4.4. Optical stimulus for model fitting

While the approaches in this study can generalize to multi-input systems (*e.g*., multiple light sources spread spatially or multiple wavelengths), only single-input systems are considered and tested here. As in Bolus *et al* ([7]), a repeated 5-second instantiation of 1 ms resolution uniform optical noise was used to stimulate spiking activity for model fitting. While the amplitude of this stimulus varied across experiments based on perceived neuronal sensitivity to light, the average range of this uniform-distributed noise was from 0 to 14.4 mW/mm^2^, and the same pattern of noise was always presented. State-space models were fit using data from the first 2.5 seconds of each stimulus trial, while the remaining 2.5 seconds of stimulation were held out and used to assess model performance.

### 2.5. Estimator

GLDS models were used both offline for designing the control law and online for estimating state feedback. For online estimation, two variants of GLDS model-based state estimation are considered. The first is a standard implementation of the Kalman filter (Section 2.5.1, [32, 33]). Another variant of this approach that was used to achieve greater robustness to plant-model mismatch was to apply Kalman filtering to estimate a parameter-augmented state vector (Section 2.5.2), which we will refer to here as a parameter-adaptive Kalman filter but has elsewhere been described as a proportional-integral (PI) Kalman filter ([34, 35]).

#### 2.5.1. Kalman filtering

The Kalman filter proceeds by alternating between a one-step prediction of the state and updating this estimate when the corresponding measurement is available ([33]). The filter has two design parameters which are reflected in the GLDS model structure (Equations 1 & 3): the covariances of the process and measurement noise, or **Q** and **R**, respectively. The value for **R** was taken from fits of the GLDS models to training data. In analyzing the performance of the Kalman filter on previously-collected spiking data, the fit matrix for **Q** was rescaled to minimize the mean squared error (MSE) between the Kalman-filter-estimated firing rate and an output of a model-free estimation method: smoothing the spikes with a 25 ms Gaussian window.

At each time point, a one-step prediction of the estimated state mean 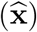, state covariance (**P**), and output 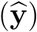 were calculated:

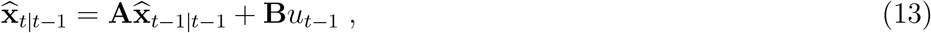

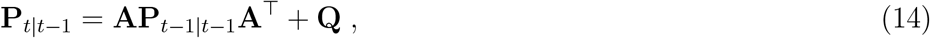

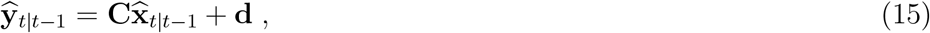

where 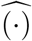 denotes estimates, (·)_*t*|*t*−1_ denotes the prediction at time *t*, given data up to time *t*− 1, and (·)_*t*|*t*_ denotes filtered estimates. Recall that all model parameters were fit to optical noise-driven spiking activity, and note that *μ* was assumed to be zero unless adaptively re-estimated (Section 2.5.2). The one-step prediction was updated taking into account the latest measurement as

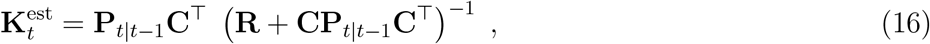

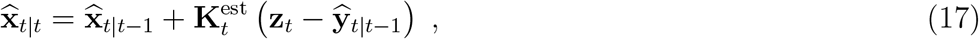

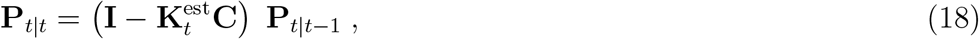

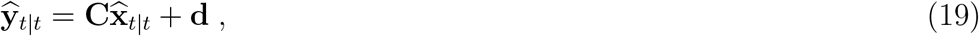

where 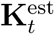 is the Kalman filter gain and **I** denotes an identity matrix.

#### 2.5.2. Parameter-adaptive Kalman filtering

For robustness of state estimation to plant-model mismatch, the state and a model parameter were jointly re-estimated by the Kalman filter. Specifically, the mean of the process disturbance, *μ*, was assumed to vary stochastically over time as a random walk:

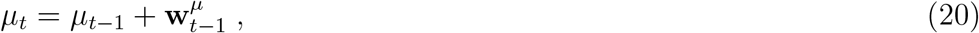

where 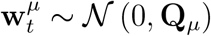 is noise disturbing the stochastic evolution of *μ*. The covariance of this process **Q**_*μ*_ effectively sets the timescale of adaptive re-estimation of *μ*. To jointly estimate this disturbance, the state and model parameters were augmented as follows:

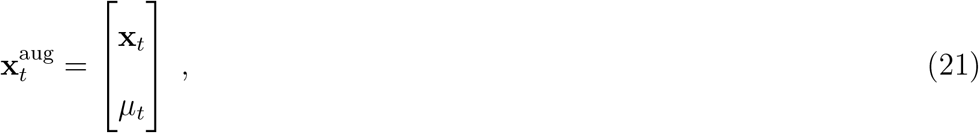

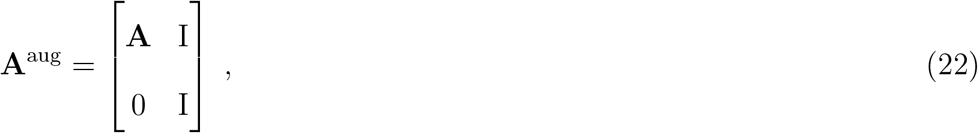

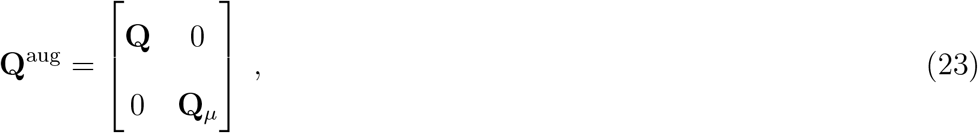

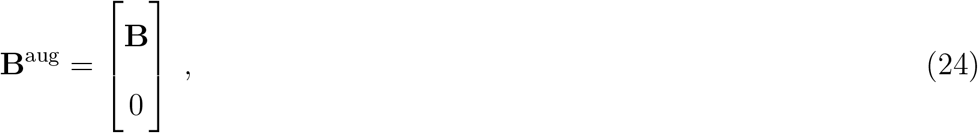

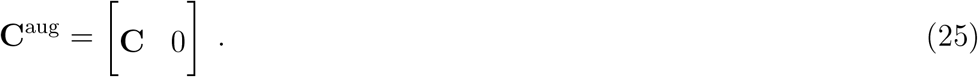

In general, such joint parameter-state estimation would require the use of the extended Kalman filter (*e.g*., [36]). However, in this case, the augmented dynamics and output equations remain linear with respect to the augmented state. Therefore, Kalman filtering was carried out on this augmented form of the state and GLDS model as detailed before in Section 2.5.1. For the purposes of this study, **Q**^aug^ was assumed to be a diagonal matrix. In analyzing the performance of this adaptive Kalman filter on spiking data, the elements of **Q**^aug^ were scaled to minimize the mean squared error between the Kalman-filter-estimated firing rate and the Gaussian smoothed estimate as before (Section 2.5.1).

### 2.6. Controller

While the state-space modeling and control framework can be readily used for trajectory tracking, the control objective in this study was holding the output neuronal firing to a fixed target, or reference, rate (*r*), corresponding to a nonzero-setpoint regulation problem ([37]), also described here as ‘clamping’.

#### 2.6.1. Control setpoint

In order to use state feedback for the case where the target is an output, we first calculated the state and optical input that would be required to achieve the target firing rate, **r**. Since this was a regulation problem, we calculated the state and input for achieving the target at steady-state. This steady-state setpoint [**y**^∗T^ **x**^∗T^]^T^ was calculated using models fit to previously collected optical noise driven data. This problem was solved by linearly-constrained least-squares [38], where the objective was to minimize the 2-norm ||**y**^∗^ − **r**||^2^, subject to the system being at steady-state **x**^∗^ = **Ax**^∗^ + **B***u*^∗^. The control signal required to achieve the target at steady state, *u*^∗^, was served as a nominal control signal, about which feedback controller gains modulated light intensity. For single-input/single-output (SISO) applications, there was a solution that resulted in zero-offset tracking (*i.e*., **y**^∗^ = **r**). However, for multi-output control where the responses to control are heterogeneous, the steady-state solutions do not result in zero-offset tracking, but rather the least-squares compromise across neurons.

#### 2.6.2. Linear quadratic regulator design

Linear quadratic optimal control was used to design controller gains *K*^ctrl^ for non-zero-set-point regulation ([37]):

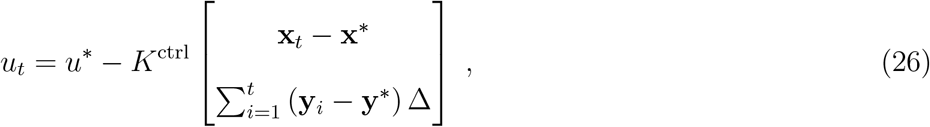

where both instantaneous state error (top row) as well as integrated output error (bottom row) were used for feedback to ensure robustness of control. Δ is the sample period (1 ms). The controller gains were chosen to minimize a quadratic cost (*J*) placed on these tracking errors and on deviations in the control ([37, 39]):

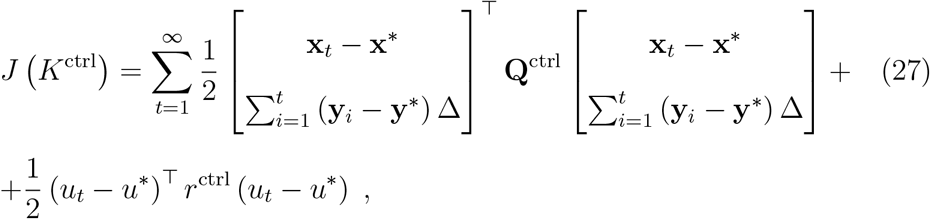

where **Q**^ctrl^ is the weight placed on minimizing squared instantaneous state error and integrated output error,

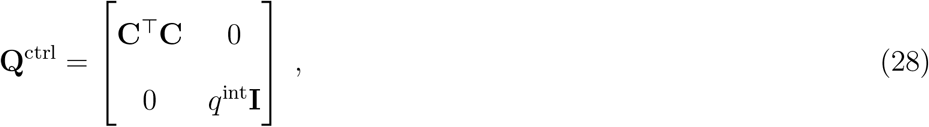

and *r*^ctrl^ is the weight placed on control deviations. Minimization of this quadratic cost function is linearly constrained by the error system dynamics

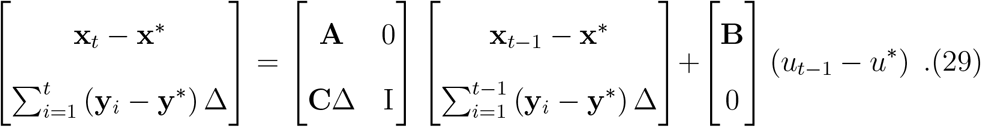

This optimization was carried out numerically by backward recursion of the discrete-time matrix Riccati equation until convergence ([37]) or calculated using the MATLAB function *dlqr()* (MathWorks). Generally, a stabilizing solution was not possible for multi-output control scenarios with integral action because of nonzero output error; however, the numerical solution for feedback controller gains still converged in practice.

#### 2.6.3. Experimental SISO control

First-order GLDS models fit to previously collected spiking responses to optical noise were used offline for designing feedback controller gains, *K*^ctrl^, (Section 2.6.2) and online for the parameter-adaptive Kalman filtering (Section 2.5.2). The diagonal elements of the assumed process noise covariance, **Q**^aug^, used in the parameter-adaptive Kalman filter ranged from 1 × 10^−9^ to 5 × 10^−8^. For controller design, the quadratic weight chosen for integral error (*q*^int^) was 1 × 10^2^, while the weight placed on control deviation, *r*^ctrl^, ranged from 1 × 10^−4^ to 1 × 10^−3^. The online-sorted spiking data fed back to the controller was used to assess performance of the control scheme; however, in cases where a 32-channel electrode array was used for recording, offline-sorted population activity was inspected to understand the local effects of closing the loop around a given putative single neuron.

#### 2.6.4. Simulated SIMO vs. SISO control

In addition to experimental validation in the SISO case, the state-space modeling, estimation, and control methods were also applied to a simulated multi-output control problem in which the objective was to push the outputs toward a common target firing rate. In this case, 5^th^-order models were used. When fitting GLDS models to SIMO datasets, we found that there was often great heterogeneity in input-output gain across outputs. Therefore, a two-output PLDS model was the simulated system being controlled, whose second output (‘neuron 2’) was a gain-modulated version of the first (‘neuron 1’), before exponentiation and spike generation. The dynamics and the first output channel of this PLDS came from a fit to an example SISO dataset. The log-linear gain of neuron 2 was swept between 0.1 and 3 times that of neuron 1. A multi-output controller and estimator were designed using a 2-output GLDS model fit to simulated PLDS data, where optical noise stimulated the PLDS in the case where the both neurons had the same gain. The neuron-averaged mean squared error performance of the SIMO control loop was compared to the SISO scenario when only neuron 1 data was fed back. For both SIMO and SISO control loops, the diagonal elements of the process noise covariance for the parameter-adaptive Kalman filter (**Q**^aug^) were all taken as 1 × 10^−6^, while the weights placed on quadratic cost of integrated tracking error (*q*^int^) versus control deviation (*r*^ctrl^) were 1 × 10^2^ and 1 × 10^−3^, respectively.

### 2.7. Performance measures

Various measures of performance are used throughout this study to quantify goodness of fit for state-space models and the effectiveness of the estimators as well as the controller.

#### 2.7.1. Model performance

The performance of GLDS and PLDS models were assessed using variance of the raw 1 ms binned PSTH explained in the held-out second half of each 5 second trial of optical noise stimulation. The variance explained was either taken as a proportion of the variance in the PSTH (pVE), or relative to the amount of ‘signal’ or explainable variance in the PSTH (pSVE, [27]). These two metrics were computed for each SISO and SIMO dataset for 5^th^ order PLDS models and 1^st^ and 5^th^ order GLDS models.

#### 2.7.2. Estimator performance

Because the control objective in this study was to track a constant reference firing rate, it was important that the estimator achieve low bias; otherwise, the integral action of the controller cannot serve its ideal purpose to eliminate steady state tracking errors. Therefore, the performance metric considered here for the online estimator was the squared bias of the single-trial-estimated firing rate compared to the corresponding spiking responses to 5-second step inputs of light.

#### 2.7.3. Control performance

To assess controller performance, the mean squared error (MSE) as well as squared bias between the achieved single-trial firing rate and the reference firing rate were calculated. Single-trial firing rate was taken as the online-sorted spike train fed back to the controller, smoothed offline with a 25 ms standard deviation Gaussian window. While MSE takes into account variance, we separately considered across-trial variability using the Fano factor ([40]) of spike counts in a 500 ms sliding window, a mean-normalized measure of spike count variability. Finally, in cases where a 32-channel multielectrode array was used for recording local population activity, the degree of synchrony between simultaneously recorded neurons was quantified in a manner similar to Wang *et al* ([41]). Briefly, a cross-correlogram was constructed by binning the relative spike times of simultaneously recorded neuron pairs. To quantify degree of synchrony, the number of correlated events in a *±*7.5 ms window (*N*_cc_) was normalized by the total number of spikes in a *±*50 ms window (*N*_tot_):

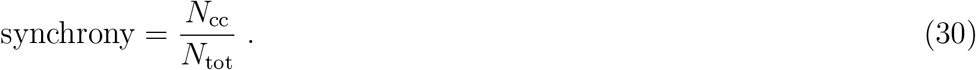

Allowing 1-second for non-steady state performance, all four of these performance metrics were calculated in a 4-second period of time during closed-loop control. As a point of comparison, the same metrics were also calculated using 4-second periods of spontaneous data recorded between trials of closed-loop stimulation.

## 3. Results

In this study, we applied a model-based optimal control framework to the experimental control of neural activity in vivo using optogenetic stimulation. Specifically, we utilized the ventral posteromedial (VPm) region of the sensory thalamus in the vibrissa/whisker pathway of the awake mouse as an experimental model system, where single-unit electrophysiological recordings were obtained while optically stimulating light sensitive channels with an inserted optical fiber. The optimal control framework relies on a state-space representation of the optically-driven dynamics of neural activity. This model is used both for the offline design of the optimal controller and for the online estimation of state feedback. Although experimental results are presented from this specific pathway and brain region, the approach is directly applicable to others. Furthermore, while the methods used here should generalize to future multi-input and multi-output (MIMO) applications, we first focus on the single-input and single-output (SISO) case where the measured outputs were single-unit spiking activity and the control objective was to track step commands (*i.e*., clamp neural activity at a fixed target firing rate). In the context of these experiments, we were able to use a linear and Gaussian model to approximate light-driven spiking responses for the purposes of controlling firing rate; moreover, we found that a low order approximation of the neural dynamics was sufficient at least for the slow timescale control/estimation objectives studied here. In experiments where multi-electrode arrays were employed to record thalamic activity, we found that simultaneously-recorded neurons responded to optical stimulation with a high degree of diversity, motivating investigation of applicability of this control framework to multi-output scenarios. We applied this framework in a simulated single-input/multi-output (SIMO) scenario, where the “output” consisted of the activity of multiple simultaneously-recorded neurons, and the control objective was to force the population activity as close as possible to a common target firing rate. Feeding back multi-output population activity to the controller enhanced the robustness of the control scheme’s ability to drive the collective population activity to a desired target in the face of heterogeneity in sensitivity to light.

Figure 1(a) illustrates the control scheme that was implemented experimentally in the awake, head-fixed mouse, where an ‘optrode’ consisting of an electrode attached to an optical fiber was inserted into the VPm. Given binned single-unit spiking activity, control and estimation was carried in realtime at 1 ms resolution using custom-written software (Section 2.2). We designed an estimator that generated an online estimate of the state of neural activity, and a feedback controller that maintained a target firing rate in the face of potential disturbances, such as reafferent sensory input (*i.e*., whisker motion) and changing brain states.

**Figure 1:**
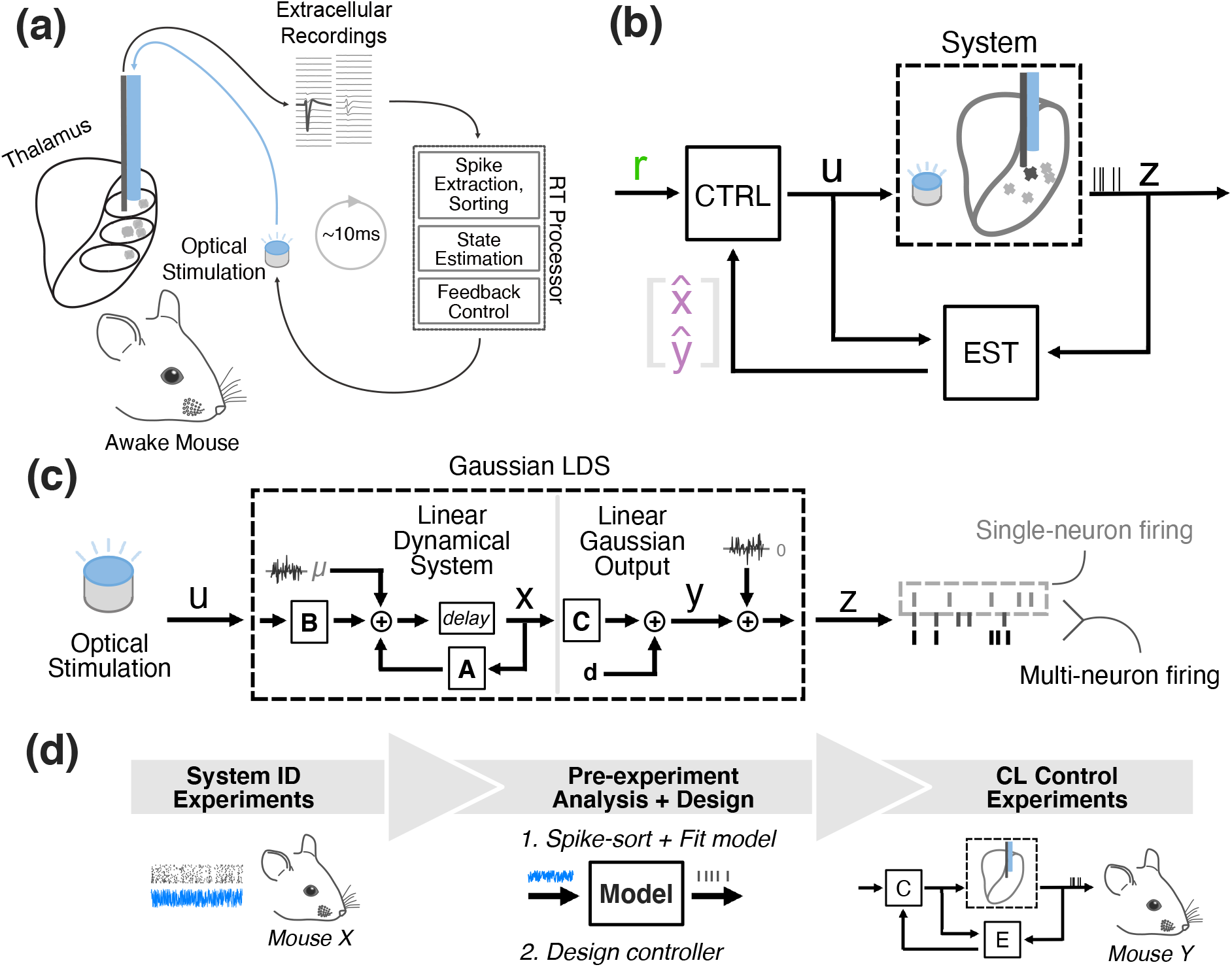
Closed-loop optogenetic control using state-space linear dynamical systems models. (a) Experimental Setup. (b) Control system block flow diagram. Spiking activity is fed back to a model-based estimator (‘EST’), which provides online estimates of the underlying state of the system (**x**) and the output (**y**), which is firing rate in the current application. The controller (‘CTRL’) uses a model to generate the system setpoint [**y**^∗T^ **x**^∗T^ *u*^∗ T^] ^T^ that corresponds to user-specified reference firing rate (**r**). An updated control signal is generated using feedback controller gains and the error between this setpoint and the online estimates of the system state/output. The updated control signal is sent to an LED driver to modulate light intensity. (c) Structure of the Gaussian LDS Model. The GLDS used throughout the control loop consists of a linear dynamical system (LDS) describing the evolution of the state (**x**) and a linear remapping of **x** to the output firing rate and eventually measured spiking (**z**). This model is used for single-neuron and multi-neuron estimation/control. (d) Workflow for closed-loop experiments. Neuronal responses to optical noise recorded in previous experiments (left) were used to fit state-space models and design the control system (middle). The resulting model-based control system was used in subsequent CL control experiments (right).

To develop a generalizable control methodology, we applied a state-space model-based control and estimation scheme where the model is used not only in the design phase but as an online estimator for the control scheme (Figure 1(b)). The model structure utilized here was a linear dynamical system (LDS), where optical input(s) modulate the activity of latent state variables. More specifically, for the purposes of this study we employed a Gaussian linear dynamical system (GLDS), in which a linear combination of the states is observed after being corrupted by additive Gaussian noise (Figure 1(c)). Here, the output of the model was either single or multi-neuron firing rate, although in principle these same techniques could be applied to other neural signals of interest such as local field potential or voltage/calcium signals. Figure 1(d) illustrates the workflow for the closed-loop experiments. Neuronal responses to optical noise recorded in previous experiments were used to fit state-space models and the control system, utilized in subsequent closed-loop control experiments to be presented in detail in later sections, highlighting the generalizability of the approach across animals.

### 3.1. GLDS captures optical noise-driven responses

The control framework used here depends on a model of the underlying dynamics for both the design of the controller and online state estimation to execute the control strategy. As we have previously described in a simpler, classical control framework ([7]), feedback control is robust to a degree of model inaccuracy. Therefore, there is an application-specific balance to be struck between model complexity/fidelity and simplicity. Here, we first asked to what extent a GLDS model could predict the experimentally observed SISO firing rate modulation with optogenetic stimulation, as this would provide a relatively simple modeling framework that is attractive in terms of its widespread applicability and ease of implementation. Since the measurements were spike counts in 1 ms bins at relatively low firing rates, a Gaussian observation model is an obvious violation of these statistics. For comparison, we also fit an LDS model whose observation model is Poisson (PLDS), which has been utilized in a range of studies for describing the dynamics of spiking neurons (Figure 2(a)).

**Figure 2:**
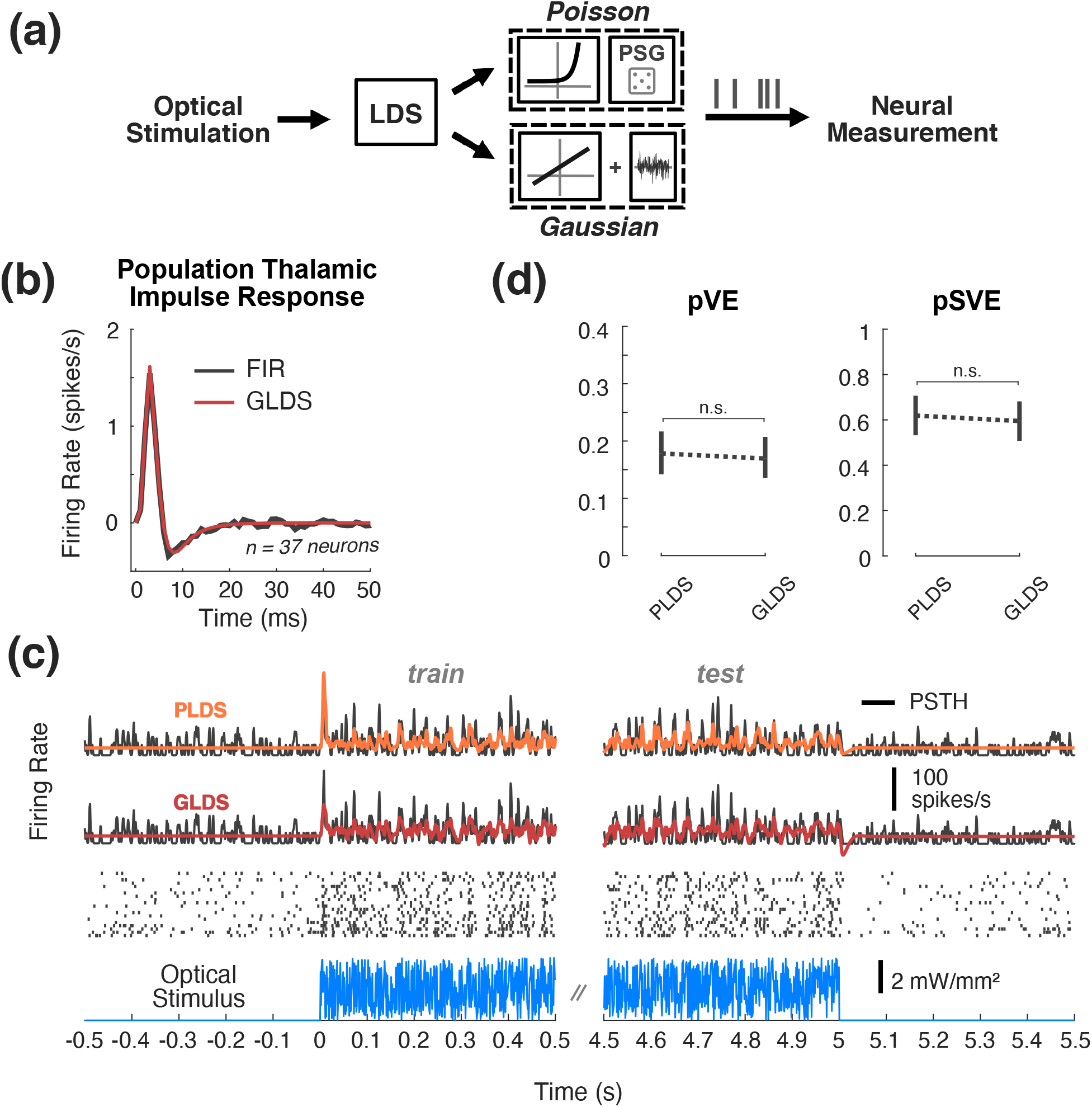
State-space models of SISO optogenetic responses. (a) SISO LDS model structure: Poisson (top) or Gaussian (bottom) output functions being considered. (b) Population impulse response. This impulse response was fit using pooled data from 37 neurons that were excited by optical noise. An FIR model fit to population data (black) is plotted alongside the impulse response from the 5^th^order GLDS model fit to the same data (red). (c) Example Data and Model Fits. Top, the PSTH (black) was smoothed with a 1 ms standard deviation Gaussian window for visualization. The fit types include 5^th^-order PLDS (orange), 5^th^-order GLDS (red). Middle, the corresponding trial-by-trial spike raster. Bottom, repeated instantiation of uniform optical noise. (d) Proportion variance in PSTH explained (pVE) and signal variance explained (pSVE) by model response to noise. All models were trained on data from first half of each trial, while model performance metrics (pVE, pSVE) were calculated from the second half of each trial. Error bars represent bootstrapped 95% confidence intervals about the population mean (n=48 neurons, 17 recordings, 9 animals).

In fitting a state-space model, the order of the model (the dimensionality of the latent state vector) must be specified. To ascertain the appropriate order of these models, we pooled together noise-driven response data from 37 neurons that were all significantly excited by the optical stimulus, and fit GLDS models to this population. For comparison, we separately fit a finite impulse response (FIR) model to the same population dataset (see Section 2.4), as it is widely used in the neuroscience literature ([31, 42]). Models were fit from recorded responses to white-noise optical inputs (Section 2.4.4). Shown in Figure 2(b) are the impulse responses for both the GLDS (red) and FIR (black) models, as a head-to-head comparison. This can be interpreted as the model prediction of the instantaneous firing rate in response to a light impulse input at time zero. Prominent in both is an initial peak at approximately 3 ms reflecting a relatively short latency excitation, followed by a subsequent drop below baseline at 7-8 ms reflecting a post-excitatory inhibition. We found that a 4^th^ to 5^th^ order state-space model was sufficient for these data, striking a balance between goodness of fit and model complexity. Note that the above analysis was restricted to thalamic neurons that were found to be excited by the optical input, which excluded other thalamic neurons that exhibited more heterogeneous behaviour (i.e. a minority of recorded neurons were indirectly inhibited by the optical input, interestingly). To capture the full heterogeneity of the population, therefore, we fit 5^th^order PLDS and GLDS models to each single-output dataset individually (*n* = 48 neurons, 17 recordings in 9 mice). A representative example SISO dataset is shown in Figure 2(c), where the firing rate estimates for the 5^th^order PLDS (first row, orange) and 5^th^order GLDS (second row, red) are superimposed onto the corresponding PSTH (black) at white-noise onset and offset. Qualitatively, there is little gained in using a PLDS model instead of a GLDS for this example, aside from the non-negativity of the PLDS firing rate. Across the population of units, there is no significant difference between the performance of the Poisson vs. Gaussian models (Figure 2(d), *n* = 48 neurons, *p* = 0.234, Wilcoxon signed-rank test). Specifically, the left plot of Figure 2(d) shows the proportion of the variance in the raw 1 ms PSTH explained by 5^th^order PLDS and GLDS fits (pVE). Note that a relatively low proportion of the variance in the raw PSTHs was explained, due to levels of intrinsic noise in the observed responses at fine timescales. For this reason, we assessed the quality of the model using a metric that takes into account the fact that some of the observed variability is not explainable across trials ([27]), instead quantifying the amount of explainable, or ‘signal’, variance the model captures. The right panel of Figure 2(d) presents the proportion of the signal variance explained (pSVE), showing that the models captured approximately 60% of the explainable variance and that there was not a significant difference in the predictive capabilities between the GLDS and the PLDS models in this dataset. Therefore, with the exception of multi-output modeling where the same PLDS versus GLDS analysis was conducted for comparison, GLDS models are used for the remainder of this study in order to leverage linear controls approaches.

### 3.2. Parameter-adaptive Kalman filtering provides robust online estimation

These GLDS models are used online as part of the Kalman-filter-based estimator (Figure 3(a), grey box) which is used to provide state feedback to the controller. While the models performed relatively well in the case of uniform white-noise optical stimulation as shown in Figure 2, when challenged with step changes in input that are often utilized in control scenarios, non-zero-mean model mismatch is clearly revealed (Figure 3(b)). In this example the open-loop model predicted firing rate (OL Prediction, red) initially under-estimates the experimentally-measured firing rate (PSTH, black) during the first second of stimulation and then consistently underestimates the firing rate at steady state. Model-based control and estimation schemes are particularly sensitive to such plant-model mismatch, as is apparent here when standard Kalman filtering used for online estimation is applied to these datasets for step changes in input. In this example in Figure 3(b) there is still an obvious bias in the average Kalman-filter estimated firing rate (KF Estimate, purple) when compared to the smoothed PSTH (PSTH, black). Moreover, because of the rapid time-course of the fit neuronal dynamics (Figure 2(b)) and the spiking nature of the measurements, the single-trial KF estimates of firing rate which will be fed back to a controller are full of extreme transients each time a new spike is measured (Figure 3(c), purple trace). Online estimation of firing rate can be made more robust by assuming there is an unmeasured, non-zero-mean disturbance that varies stochastically (e.g., other exogenous inputs), augmenting the state with the mean(s) of this disturbance (*μ*), and jointly re-estimating this along with the state using Kalman filtering (Figure 3(d), see methods for details), which we refer to here as the parameter-adaptive Kalman filter, but has elsewhere been described as a proportional-integral Kalman filter ([34, 35]). As can be seen in the example in Figure 3(e), this adaptive Kalman filter produces an effectively unbiased estimate of the experimentally-observed PSTH in SISO applications (Figure 3(e), purple vs. black), and it is able to do so with a single-trial estimate of firing rate that is smoother than that achieved by the standard Kalman filter (Figure 3(f), *cf*. Figure 3(c)). In this example, the parameter-adaptive Kalman filter approach accounts for apparent model mismatch by estimating a process disturbance *μ* that on average pushes the firing rate above the model prediction for the first second of optical stimulation and then pulls the estimated firing rate below that prediction at steady state (Figure 3(g)). The filtering approach works well in this illustrative example and at a population level, as it brings the estimation bias to near-zero levels compared to the standard Kalman filter (Figure 3(h), *p* = 1.63 × 10^−9^, Wilcoxon signed-rank test, *n* = 48 neurons, 17 experiments, 9 animals). At least in the context of estimating step responses, we see there is little benefit in using a 5^th^order versus 1^st^-order GLDS model for this SISO application (Figure 3(h), black, *p* = 0.0830, Wilcoxon signed-rank test). Importantly, the parameter adaptation provides enough robustness that even the population-average GLDS model in Figure 2(b) was able to estimate SISO firing nearly as well as models fit to each neuron individually (Figure 3(h), gray, *p* = 0.0142, Wilcoxon signed-rank test). Since the control objective in this study is to clamp firing rate at relatively long timescales, we therefore used a 1^st^-order Gaussian approximation for the system. However, for fast timescale trajectory tracking problems, a higher-order model would almost certainly be warranted (see Discussion), and higher-order models are important even for long timescale control/estimation in multi-output scenarios (see Section 3.6).

**Figure 3:**
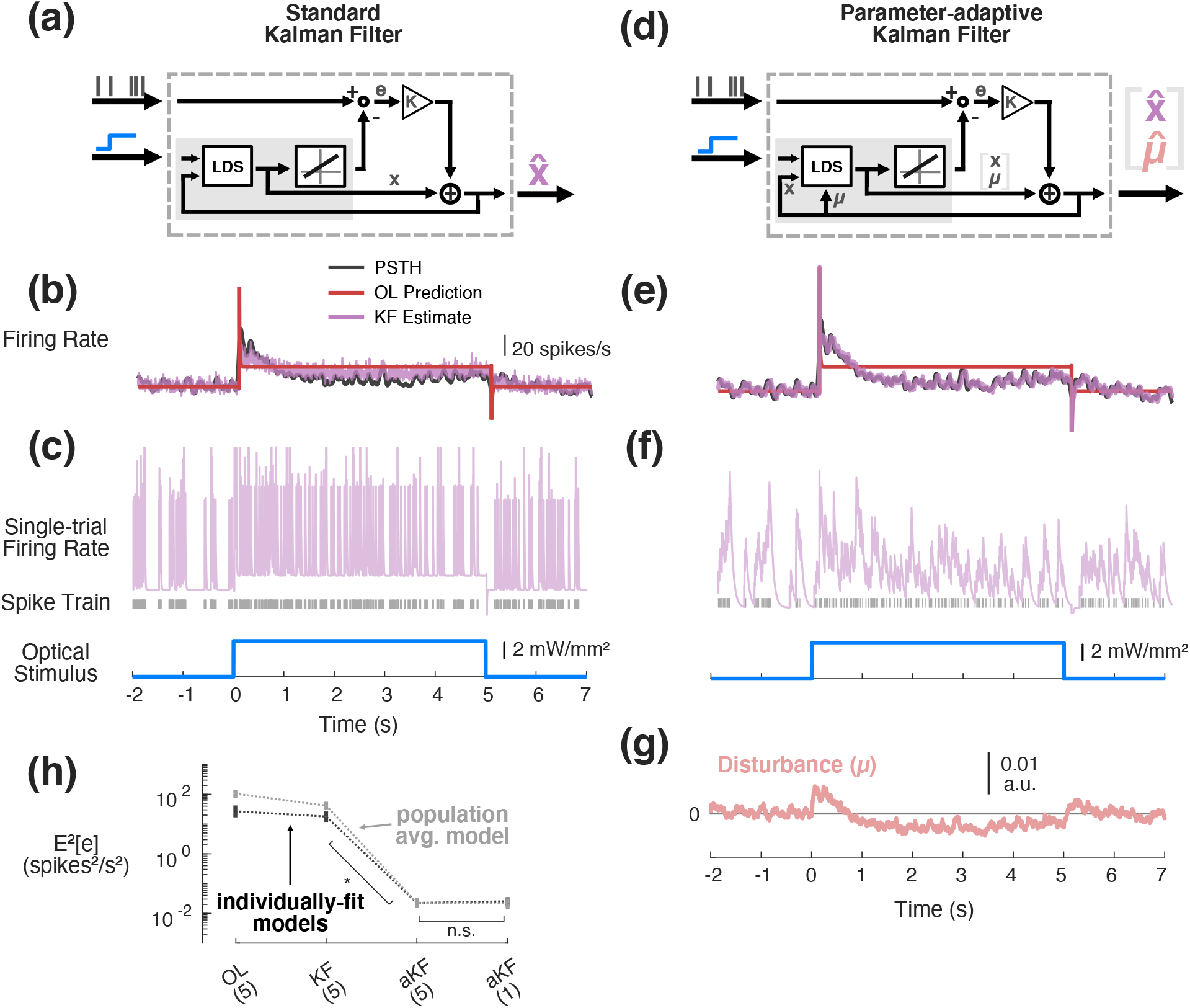
Kalman filtering for online estimation in SISO applications. (a) Standard Kalman filter. Prediction error (*e*) is used to correct the estimate of state at each time step. (b) Example open-loop (OL) prediction of neuronal response (red) to step input of light (blue) using 5^th^order GLDS fit to noise-driven data, compared to PSTH smoothed with 25 ms Gaussian window (black) and the trial-averaged response estimated using the standard implementation of the Kalman filter (5^th^-order GLDS) (purple). (c) Example single-trial Kalman filter estimate (purple) along with corresponding spike raster (grey). (d) Parameter-adaptive Kalman filter. In addition to estimating the state of the system, this approach jointly re-estimates a state disturbance (*μ*) at each time step. (e) Same as (b) but trial-averaged estimate of firing rate using the parameter-adaptive Kalman filter. (f) Same as (c) except single-trial estimate using parameter-adaptive Kalman filter. (g) Trial-averaged disturbance on the first state estimated using parameter-adaptive Kalman filter. (h) Population average squared-bias in estimation calculated between the single-trial spiking responses and the OL prediction of a 5^th^-order GLDS, the standard Kalman filter using the 5^th^-order GLDS, and the parameter-adaptive Kalman filter (aKF) using a 1^st^- or 5^th^-order GLDS. Black and grey data points correspond to error associated with using individually-fit models vs. a single population average fit model, respectively. Error bars represent bootstrapped 95% confidence intervals about the mean (n=48 neurons, 17 recordings, 9 animals).

### 3.3. State-space control performs well in SISO clamping applications

The control and estimation framework was tested experimentally in the awake head-fixed mouse in a SISO configuration, where spiking activity of a single neuron was fed back to a controller with a single channel of optical input. The models used for pre-experiment controller design and for the online estimation for state feedback were fit to thalamic spiking responses to optical noise from previous experiments in separate mice, as illustrated in Figure 1(d). As previously noted, 1^st^-order GLDS models were sufficient and were therefore used for this particular application; however, higher order models would be merited or necessary in other scenarios. The robustness of the estimator (Figure 3), the use of feedback, and the slow timescale nature of the control objective allowed GLDS models fit to previously-collected noise response data to be used for experimental control and estimation, rather than fitting a model during an experiment, the timespan of which is limited in the context of awake, head-fixed recordings. The feedback controller was designed using output-weighted LQR ([37]), where the state of the system was augmented with the integrated output in order to find not only proportional feedback gains on the state, but integral feedback gains to minimize steady state tracking errors (Section 2.6.2). Additionally, since this particular application is a non-zero setpoint regulation problem, the steady-state set-point of the system [**y**^∗ T^ **x**^∗ T^ *u*^∗ T^] ^T^ at the desired output firing rate (*r*) was calculated as described in Section 2.6.1.

Figure 4 illustrates the performance of the control framework for a typical single thalamic neuron and the summary performance across experiments. Figure 4(a) is an illustration of the control implementation, highlighting the feedback controller and the online estimator. In the case of the estimator, Parameter-adaptive Kalman filtering is being used to estimate not only the state of the system being controlled but also the uncontrolled disturbance (Figure 4(a), estimator block). On the other hand, the controller is operating on the error between the estimated state of the system and the desired steady-state set point as well as the integrated output error (Figure 4(a), controller block). In this example (Figure 4(b)), the baseline ongoing activity of the recorded neuron was approximately 5 spikes/s, and the controller was activated at time zero with a target firing rate of 20 spikes/s. Upon activating the controller, the neuron reached and remained at the target firing rate (green), as reflected in the average firing rate (black). Importantly, the controller operated using online estimates of state and corresponding output firing rate provided by the estimator (Figure 4(b), purple). The firing rate of the online estimator (purple) also quickly reached the target (green) and remained there. As shown previously in Figure 3, the online estimate was on average unbiased, as it matched the offline estimate of the average firing (black, PSTH smoothed with 25 ms s.d. Gaussian). The controller achieved the target with well-below spontaneous levels of across-trial variability, quantified using the Fano factor (FF) that captures the spike count variance relative to the mean spike count (Figure 4(b), middle). In this particular example, the controller’s use of feedback resulted in a gradual increase in light intensity that was needed to maintain the target level of spiking over the control epoch. Also note that this control signal varied substantially across individual trials (Figure 4(b), bottom, light blue), with significant individual trial variability serving to drive the firing rate tracking and quench the variability. While variable across experiments, in general it was approximately 1.1 seconds before the controller was able to push neuronal firing to within 2% of its steady state value (bootstrapped 95% confidence intervals about the median, 0.47 to 1.34 sec, n=11 recordings). This settling time metric was calculated by fitting a second order transfer function to closed-loop step data and using the MATLAB ‘stepinfo’ function (Mathworks, Inc.).

**Figure 4:**
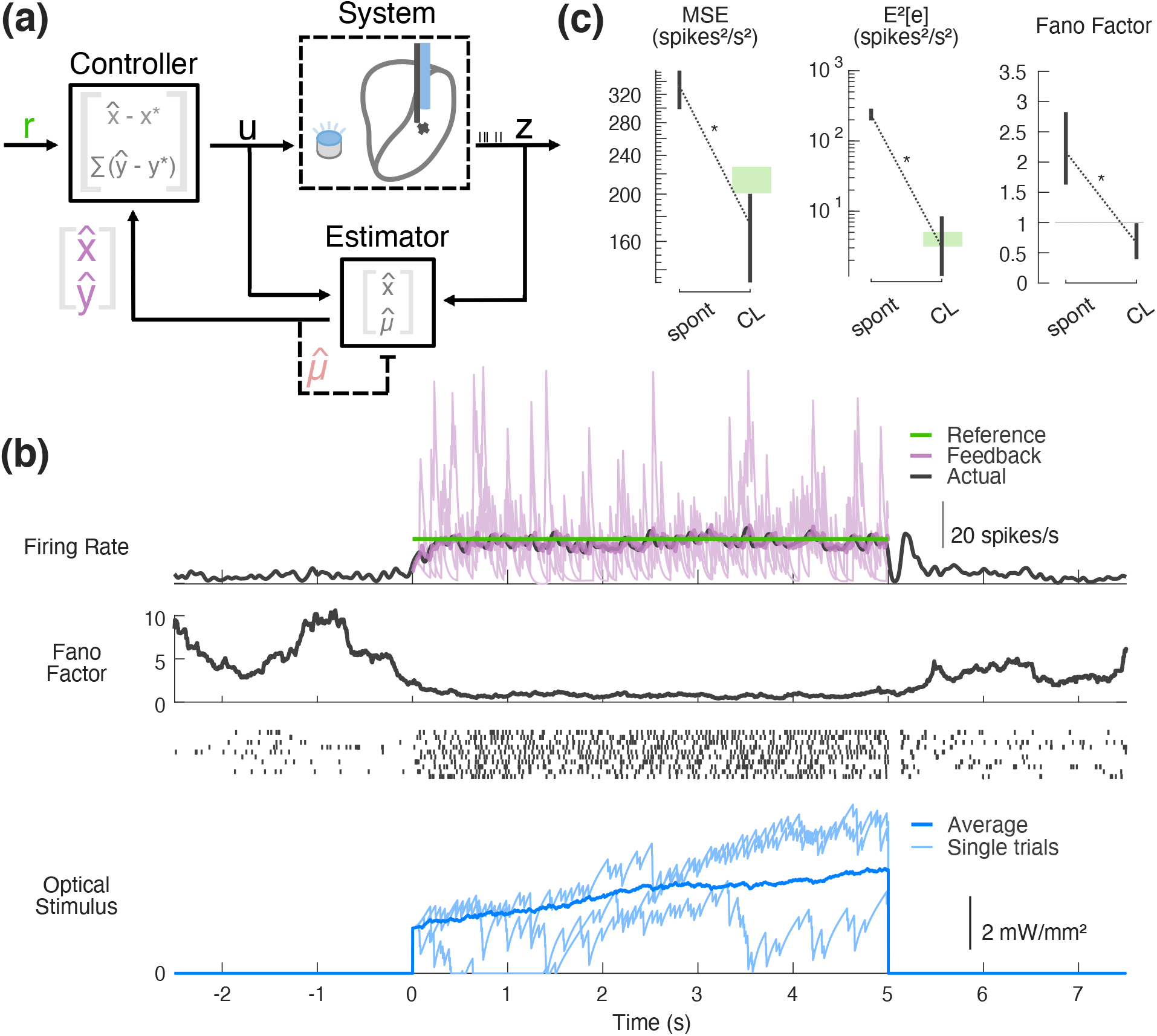
Experimental SISO control and estimation. (a) SISO control block flow diagram. Shown inside the controller and estimator blocks are the notions of state being used in each operation. (b) Example experimental SISO control. (top) Fed-back online estimate in purple (single trial in light purple, trial-averaged in bold), along with the corresponding trial-average offline estimate (25 ms s.d. Gaussian-smoothed PSTH); (middle) across-trial spike count variability (Fano factor in 500 ms sliding window) and corresponding example spike rasters from 10 randomly selected trials; (bottom) controller input. (c) Population controller performance. In spontaneous vs. closed-loop (CL) control conditions, mean squared error (left) and squared bias (middle) were calculated between the reference (20 spikes/s) and single-trial feedback spiking data smoothed with a 25 ms s.d. Gaussian window; average Fano factor was also calculated (right). For each trial, four seconds of spontaneous data were compared to four seconds of CL control data. The first second was ignored in order to obtain a measure of steady-state performance. Error bars represent bootstrapped 95% confidence intervals about the mean. Green bands represent 95% confidence band for the metrics calculated from simulated Poisson firing at the target rate.

Across experiments (*n* = 11 neurons, 11 experiments), the control framework performed well as quantified by the summary of performance metrics in Figure 4(c). For each of these metrics, the measure during closed-loop (CL) control is compared to that from the spontaneous period (spont) before the control was activated at time zero. The mean squared-error (MSE) between an offline estimate of the single-trial firing rate and the target (Figure 4(c) left) decreased significantly with activation of the control law as expected (*p* = 0.00195, Wilcoxon signed-rank test), and the MSE during closed-loop control was even below that of a Poisson spike generator driven at the target rate (green bar), consistent with the sub-Poisson variability as revealed by the Fano-factor in Figure 4(b). Because the MSE captures a combination of the variance and the bias, we separately computed the bias in the control (Figure 4(c) middle), substantially reduced with the activation of the control (*p* = 0.000977, Wilcoxon signed-rank test) and at the level expected for a Poisson spike generator driven at the target rate (green band). To further quantify the reduction in across-trial variability during the control, we computed the average Fano-factor in a 500 ms sliding window, exhibiting substantial reduction from supra-Poisson variability (FF*>*1) in the spontaneous activity to sub-Poisson variability (FF*<*1) during the control (Figure 4(c) right).

### 3.4. Multi-electrode recordings reveal effects of SISO control on simultaneously recorded neurons

Up to this point, the state-space control framework has been shown effective for tracking step commands in single-neuron scenarios. However, neural recording methodologies (electrophysiology and imaging) continue to scale in size (*e.g*., larger numbers of channels for electrophysiology, or pixels for imaging) and one of the main benefits of using state-space models for control and estimation is the generalizability to such multi-output problems. While the preceding experimental demonstration was presented in the context of a single channel of light input and a single channel of neuronal output, in a subset of experiments, we simultaneously recorded multiple nearby neurons in the thalamus of the awake, head-fixed mouse. This provides a window into the effect of the stimulation on the local population while a single neuron is used as an ‘antenna’ around which the controller is operating, which we will refer to as the feedback (FB) neuron (Figure 5(a)). For the purposes of this analysis, we inspected simultaneously-recorded neurons that were excited by 5 ms square pulses of light with sub-10 ms latency. Figure 5(b) provides an example in which one neuron is being used for feedback (purple, top), while offline spike sorting reveals the activity of six other simultaneously recorded neurons, which we will refer to as non-FB neurons (black, trial-averaged firing rates; green shows control target). While nearby on this 25 *μ*m spaced electrode array (Figure 5(b), right), these neurons nevertheless responded heterogeneously to the optical stimulation. In this particular example, the FB neuron was substantially more sensitive to light compared to the non-FB neurons, as evidenced by their modest response following the controller activation at time zero. While all increase their firing rates in response to the controller input, none are driven to or above the target firing rate of 20 spikes/sec in this example. This was not always the case, as in other experiments the non-FB neurons could be either more or less sensitive to the light input as compared to the FB neuron. Across experiments (*n* = 8 feedback neurons, 23 non-feedback neurons, 8 experiments), we calculated the average per-trial firing rate during the pre-control spontaneous (spont) versus control periods for the FB neuron and the non-FB neurons recorded simultaneously. As expected, for the FB neurons, the controller reliably pushed the firing rate to the 20 spikes/s target. In contrast, while the average firing rate of non-FB neurons was significantly elevated from spontaneous levels and toward the 20 spikes/s target (*p* = 0.00781, Wilcoxon signed-rank test), it did so with high variability as evidenced by very wide confidence intervals about the across-experiment average (12.7 to 25.1 spikes/s, Figure 5(c), left, grey) and made more plain by the fact that FF did not change from its spontaneous levels in the non-FB neuron case (Figure 5(c), right, grey, *p* = 0.844, Wilcoxon signed rank test).

**Figure 5:**
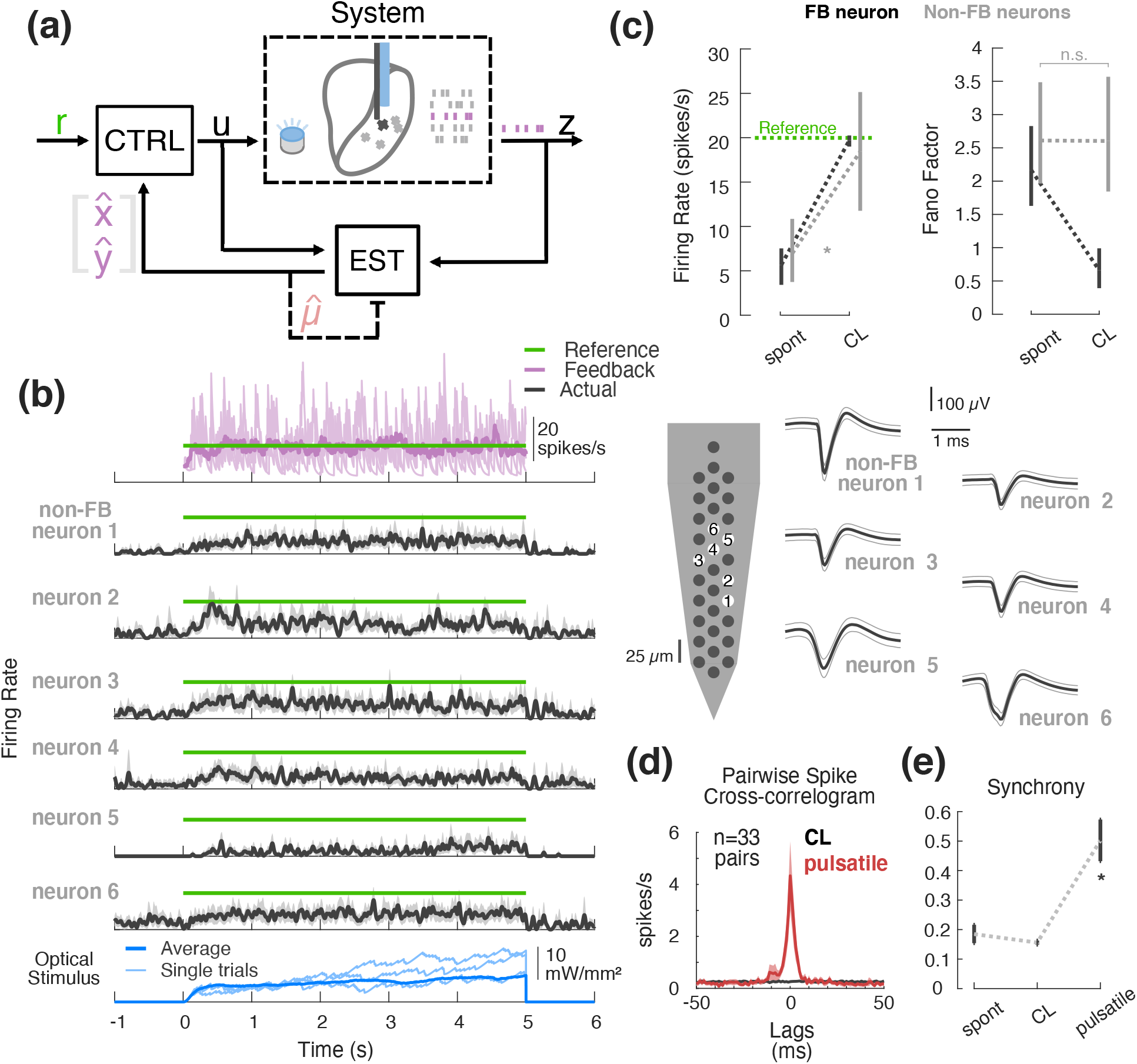
Effects of SISO control on local population. (a) SISO control block flow diagram with multi-output recordings. (b) Example experimental SISO control with simultaneous multi-output recordings. (top) Fed-back online firing rate estimate in purple (single trial in light purple, trial-averaged in bold) relative to reference (green); (middle) trial-averaged firing rate estimates for simultaneously recorded non-FB neurons (25 ms s.d. Gaussian-smoothed PSTH); (bottom) controller input. To the right are the waveforms of each neuron in this example (average waveform in black, *±*1 s.d. in grey). (c) Spontaneous vs. CL population average firing rate (left) and Fano factor (right) for the feedback neuron (black) as compared to the other non-feedback-neurons recorded simultaneously (grey). Error bars represent bootstrapped 95% confidence intervals about the mean. (d) Population spike cross-correlogram of simultaneously recorded pairs during optical stimulation. Bold black represents population mean in each 1 ms bin for CL, while fills represent 2 standard errors about the mean. For comparison, red represents population average spike cross-correlogram for response of the same cells to 5 ms square pulses of light presented in open-loop. (e) Population synchrony for spontaneous vs. closed-loop vs. pulsatile conditions. Synchrony was taken as the number of spikes occurring in the *±*7.5 ms bins, relative to the total number of spikes in the *±*50 ms window. Error bars represent 95% confidence intervals about the mean.

Beyond the firing rate of individual neurons within the population, it is important to determine what effect the optical stimulation has on the spike timing and synchronization across the population. Although we have previously shown that optical stimulation over some ranges results in a somewhat reduced synchronization relative to comparable electrical stimulation ([21]), it remains an important issue to quantify the effect in the context of the control scheme used here. We find that the use of continuously graded closed-loop stimulation did not significantly synchronize the recorded thalamic neurons when compared to commonly used pulsatile stimulation (Figure 5(d-e)). Spike cross-correlograms were calculated from relative spike times for each of 33 simultaneously-recorded pairs of neurons (see Section 2.7.3). The population correlogram shows no peak at or around zero-lag for the case of closed-loop control (Figure 5(d), black). In contrast, 5 ms square pulses of light delivered in open-loop at 10 Hz to the same neurons caused clearly aligned spiking (Figure 5(d), red). There was very little synchronization of recorded neurons during closed-loop control epochs compared to the results using pulsatile stimulation (Figure 5(e), *p* = 5.39×10^−7^, Wilcoxon signed-rank test), where synchrony was quantified as the number of temporally-aligned spikes in *±*7.5 ms window, relative to the total number of spikes in a *±*50 ms window. Note that these open-loop pulses were in general higher amplitude than the continuously modulated closed-loop stimulation, so it is not necessarily the case that pulsatile inputs would have such synchronizing effects at all stimulation intensities. Also, it is possible that low-amplitude pulsatile stimulation may provide a smaller amount of heating as compared to sustained light inputs used by this control strategy. That said, the estimated light levels used by the controller tended to be on the order of (or less than) 10 mW/mm^2^ which corresponds to less than 1 mW of optical power from the optic fibers used in this study. A previous study that measured the neuronal effects of optical stimulation in the absence of opsin expression reported no significant change in firing rate for 1 mW light intensity shone through a 200 μm fiber ([43]). While it is therefore unlikely the levels of stimulation used here led to heat-induced changes in neuronal activity, this is certainly an important consideration moving forward.

### 3.5. GLDS models generalize to multi-output datasets

In the previous section, we considered the effects of closed-loop optogenetic control on nearby neurons when the control was applied in a single FB neuron (*i.e*., SISO) scenario. More generally, the goal of control may be bringing the firing rate of a neuronal population toward a common target, rather than a single neuron, in order to provide a more controlled and uniform input to downstream neurons. An open question is whether multi-output control would serve this goal better than the above single-neuron ‘antenna’ approach. One of the strengths of the state-space control and estimation framework is that it is amenable to such multi-output applications.

To investigate the multi-output capabilities of this approach, we first demonstrate that these GLDS models can be used to capture the SIMO systems in cases where we recorded multiple neurons simultaneously. We found that the response of multiple neurons to optical noise could be represented by 5^th^order GLDS models due to the similarity in dynamics and coupling across the channels. Note that in contrast to the single-output scenario considered previously, simultaneously recorded neurons are taken to be output channels driven by a common LDS. In other words, a common state vector is mapped to individual outputs (Figure 6(a)). The same subspace algorithm was used to identify these multi-output models as before for the SISO case. Figure 6(b) and (c) provide example results of the GLDS state-space modeling for an example set of four thalamic neurons recorded simultaneously. Figure 6(b) shows the impulse response of the GLDS model of the dynamics across these recorded neurons (red), superimposed on the corresponding FIR estimates (black), showing good correspondence as previously exhibited for the single neuron case in Figure 2(b). Figure 6(c) shows the model predictions of the responses to uniform white noise optical stimulation (red) as compared to average experimentally recorded trial averaged firing (black) for this same set of neurons. The neurons clearly responded heterogeneously to light in terms of overall gain, and the GLDS model captures this and the temporal characteristics of the response to optical noise well. On average (*n* = 11 experiments, 42 neurons), 5^th^order GLDS models predict population PSTHs approximately as well as in the previously shown SISO case (pSVE 60%, Figure 6(d)). As before (Figure 2), multi-output PLDS models were also fit to the same data and we found no significant difference between the performance of the Poisson vs. Gaussian LDS models in explaining the PSTHs under these conditions (*p* = 0.923, Wilcoxon signed-rank test). As is clear in the example responses in Figure 6(c), across recordings we found there was often large (sometimes tenfold) heterogeneity in overall sensitivity to light as measured by the static input-output gain, even though the dynamics could be qualitatively similar. To explicitly characterize this heterogeneity, Figure 6(e) represents the static gain for each recorded neuron, calculated from the steady state input-output gain of the GLDS fits (circle represents mean, bars represent range). It should be noted that among the inclusion criteria for this study was that neurons must be significantly modulated by light (Section 2.3); however, this does not mean that all neurons were directly stimulated and so could be indirectly excited or even inhibited (*i.e*., have negative gains) by optical stimulation of ChR2 expressed in other cells in the network.

**Figure 6:**
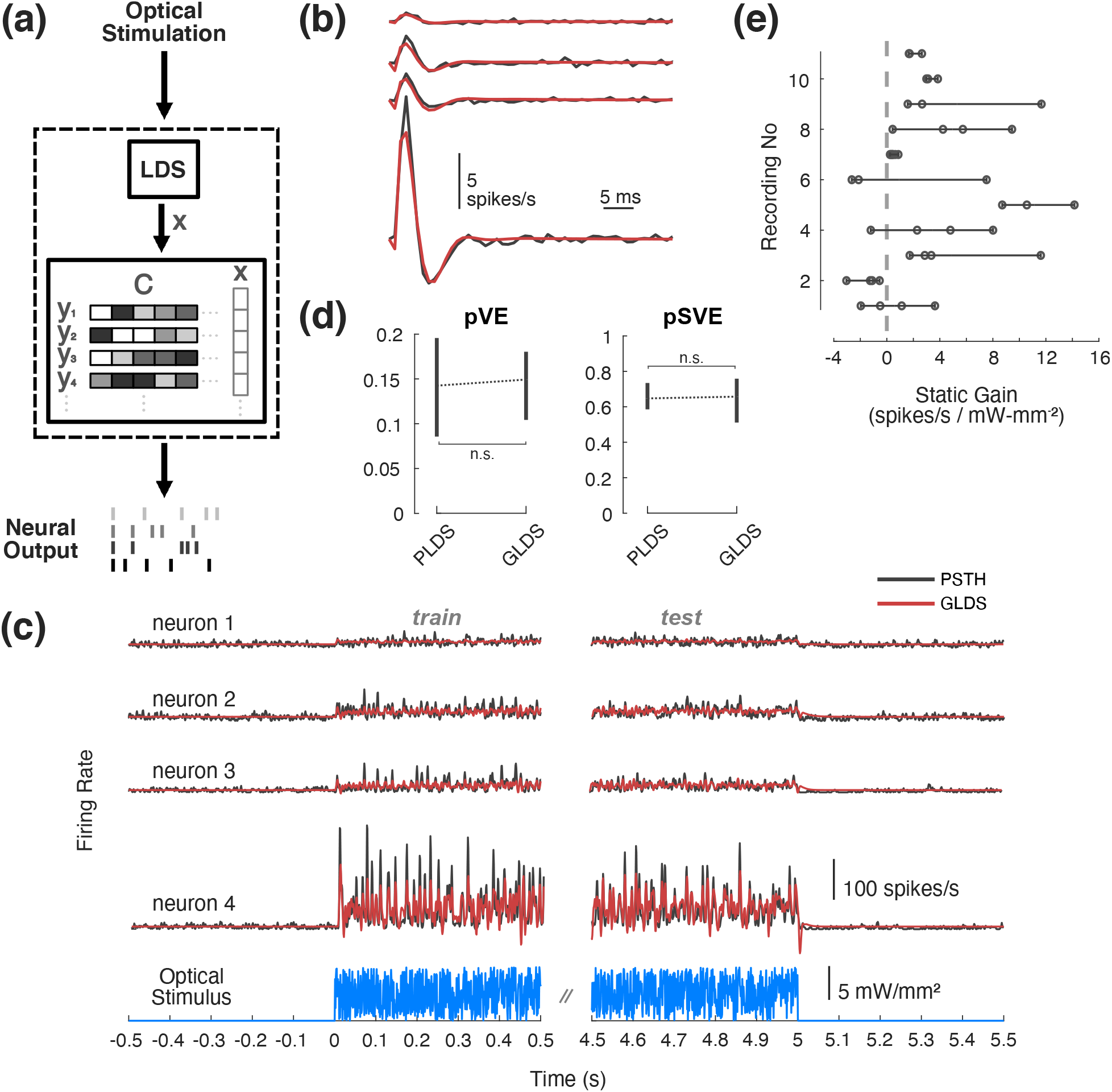
State-space models of SIMO optogenetic responses. (a) Multi-output GLDS model diagram. A common state is mapped to multiple outputs. (b) Example impulse responses from a single multi-output GLDS model (red) vs. multiple single-output FIR models (black). (c) (top) Example multi-output GLDS model response to optical noise (red) vs. PSTH (black). (bottom) Optical noise stimulus used to fit the model. (d) Population proportion variance in PSTH explained (pVE) and signal variance explained (pSVE) by model response to noise. All models were trained on data from first half of each trial, while model performance metrics (pVE, pSVE) were calculated from the second half of each trial. Error bars represent bootstrapped 95% confidence intervals about the population mean. (e) Range of static input-output gain across and within recordings.

### 3.6. Online estimation methods generalize to multi-output applications

As described previously, the state-space models were used for both offline controller design as well as for the online estimation of feedback provided to the controller. Figure 7 details the performance of the parameter-adaptive Kalman filter estimator for the single-input, multi-output (SIMO) case. While a 1^st^-order GLDS Kalman filter performed sufficiently well as an estimator for the SISO application (Figure 3), a 1^st^-order approximation leads to substantial bias in the estimates in multi-output scenarios, as shown in the example 3-neuron recording of Figure 7(a) (trial averaged estimate in dark purple vs. PSTH in black). In this example, the adaptive Kalman filter overestimates the firing activity of neuron 1, while underestimating the firing of neurons 2 and 3. Moreover, the filter fails to capture the gradual decline in firing of neuron 3 over the course of the step response. This is to be expected, as there is only one state disturbance being estimated in the 1^st^-order case for multiple outputs that may be independently perturbed. For the same multi-output example, a higher order adaptive Kalman filter (5^th^-order, Figure 7(b)) achieves substantially lower estimation bias, albeit not unbiased like the SISO scenario (Figure 3). In this example, the average activity of neurons 1 and 3 is accurately estimated; however, the firing rate of neuron 2 is under-estimated initially. Across the population of multi-output recordings (*n* = 11 experiments, 42 neurons), the 5^th^-order adaptive Kalman filter provides lower estimation bias than a standard Kalman filter (Figure 7(c), *p* = 2.37 × 10^−8^, Wilcoxon signed-rank test) as well as a 1^st^-order adaptive Kalman filter (*p* = 2.20 × 10^−8^, Wilcoxon signed-rank test). It is unsurprising that the higher order adaptive filter improved performance in the multi-output case because there are more state disturbances being estimated. However, as is evident in Figure 7(b), it is important to note that while higher order models perform better, this form of parameter-adaptive Kalman filtering does not independently minimize the estimation error for each output neuron in general because the estimated process disturbances act on a set of common state variables.

**Figure 7:**
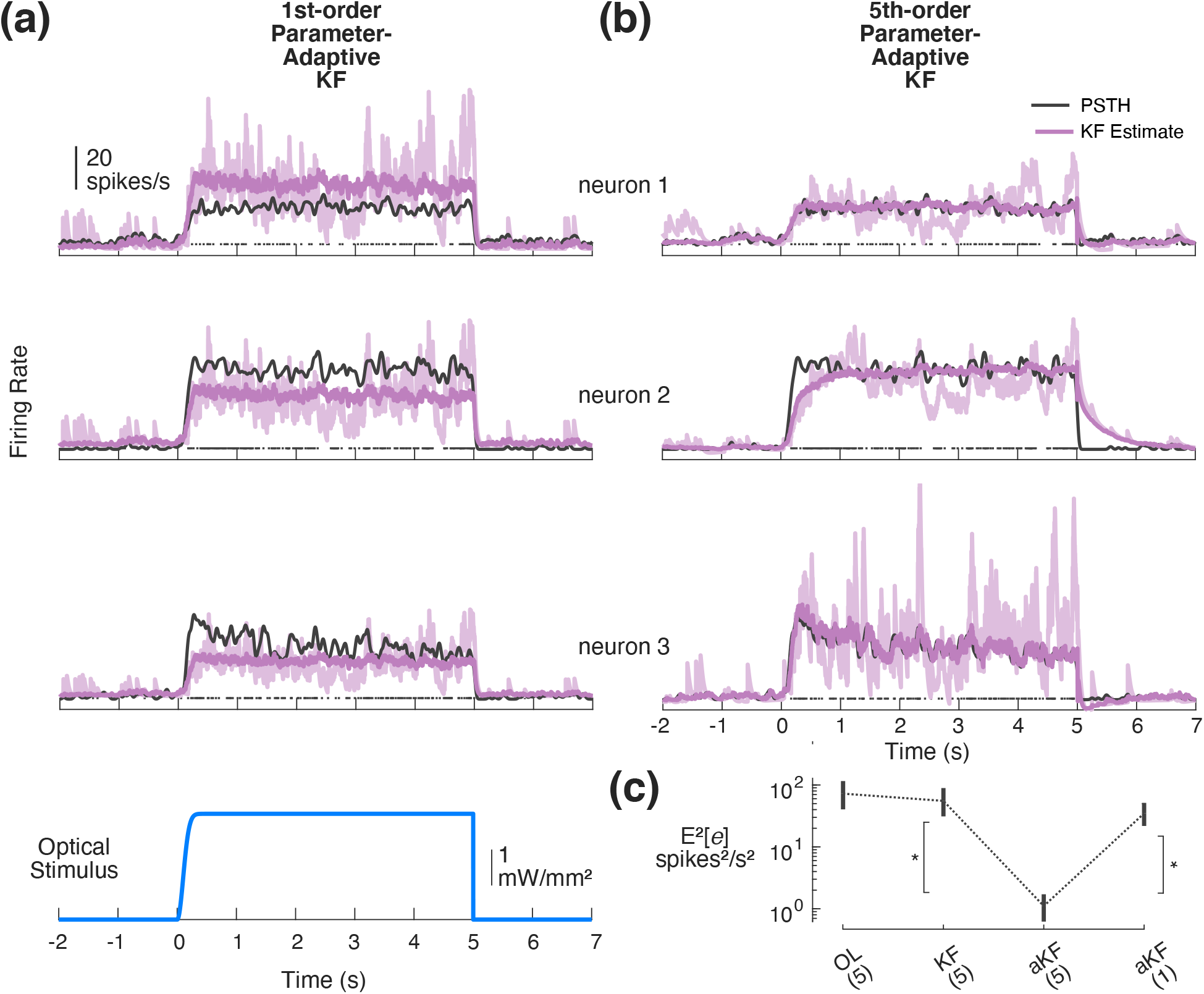
Kalman filtering for online estimation in SIMO applications. (a) Online estimation for example SIMO system using parameter-adaptive Kalman filter with a 1^st^-order GLDS. Shown for each of three simultaneously recorded single neurons: PSTH (black), trial-averaged estimate (bold purple), and example single trial (light purple). (b) Online estimation of firing rate using parameter-adaptive 5^th^-order GLDS (same data as in (a)). (c) Population summary squared-bias: Open-loop (OL) prediction of 5^th^order GLDS, standard Kalman filter (KF), and parameter-adaptive KF (aKF) for 1^st^- and 5^th^-order models. Fills/error bars represent 95% confidence intervals about the mean.

### 3.7. Simulated SIMO control more robust to population heterogeneity than a SISO ‘antenna’ approach

While we experimentally tested the control framework in the context of single-neuron feedback, the optogenetic approaches we and others use results in opsin expression in a region of tissue rather than just the feedback neuron. Furthermore, the light source used to stimulate activity impacts a volume of tissue. So, optical stimulation naturally affects a local population of neurons, each of which is potentially different in terms of intrinsic excitability, levels of opsin expression, *etc*., resulting in a net change in sensitivity to light. Because downstream neurons receive inputs from multiple cells, it is important to investigate the applicability of this technique for feedback control of a local population, rather than a single neuron. To demonstrate the generalizability of this approach to multi-output control problems and to explore the robustness of SIMO control to population heterogeneity in light sensitivity (Figures 5(b), 6), we simulated a 2-output system whose second output (neuron 2) ranged from much less sensitive than neuron 1 to much more sensitive (Figure 8(a)). To simulate spiking, an example PLDS model previously fit to SISO experimental data was chosen to most accurately represent the complexities in the data (*e.g*., spiking). The output matrix of this model was augmented with a second row whose elements were gain-modulated versions of the first row (Figure 8(a), bottom inset). This log-linear gain term was swept from 0.1 to 3 times that of neuron 1. After fitting GLDS models to simulated SIMO datasets (below), this resulted in linear gain of neuron 2 ranging from 0.027 to 13.3 times that of neuron 1. In these simulations, the dynamics of the PLDS model neurons were held fixed. This resulted in a set of simulated datasets representing a range of similarity between the two output neurons.

**Figure 8:**
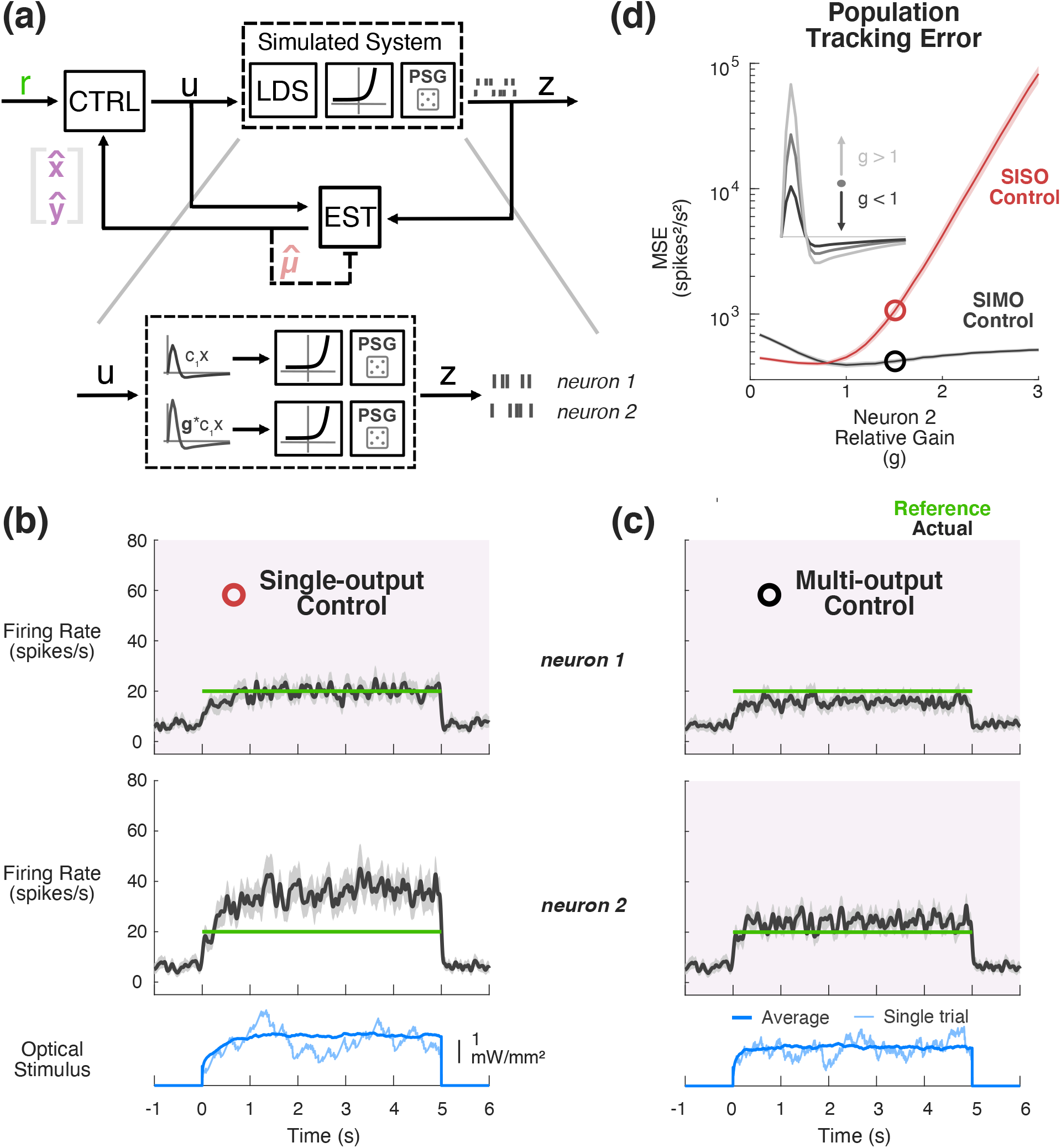
Simulated SIMO control and estimation. (a) Simulated control of a SIMO PLDS model system. A 2-output PLDS was simulated, whose second channel of output was a gain-modulated version of the first channel. (b) Example SISO Control. Only data from neuron 1 (top) was fed back during the simulated control. PLDS neuron 2 was 1.5 times more sensitive than neuron 1 before exponentiation in this example. Actual firing rate of each neuron (PSTH smoothed with 25 ms s.d. Gaussian) is shown in black as compared to reference (green). (c) Example of multi-output control. Both neurons’ data are fed back to controller for online estimation and control. Signals are the same as in (b). (d) Simulated mean-squared tracking error for SISO control of neuron 1 (red) vs. SIMO control (black) as a function of the relative log-linear gain of neuron 2. Circles denote mean-squared error for examples in (b), (c).

Using previously described methods, a multi-output GLDS model was fit to simulated spiking responses to optical white noise in the case where neurons 1 and 2 had the same log-linear gain. A single-output model was fit to the responses of neuron 1. An estimator and controller were designed using these SIMO and SISO models and they were applied to control of the 2-output PLDS across a range of output gain disparities. In the SISO case, only the activity of neuron 1 was used for feedback control, while in the SIMO, spiking activity from both neurons was fed back to the estimator and controller. Examples where the log-linear gain of neuron 2 is 1.5-times that of neuron 1 are provided in Figure 8(b) and (c), showing the SISO and SIMO control results, respectively. Qualitatively, while SISO control of neuron 1 successfully clamps activity of that channel at the target 20 spikes/sec, neuron 2 is well above the target (Figure 8b). Conversely, the multi-output control case strikes a balance between the two, allowing neuron 1 to fall below the target and reigning in the above-target activity of neuron 2 (Figure 8(c)). As summarized in Figure 8(d), population tracking performance was quantified as the total MSE of neuron 1 and neuron 2 single-trial firing rates vs. a 20 spikes/s reference firing rate, both for single-output (red) and multi-output (black) control strategies. Highlighted with the open symbols are the performances of the examples given in Figure 8(b) and (c), where the value of the log-linear gain of neuron 2 was 1.5 times that of neuron 1. As expected, a multi-output control strategy is more robust to population heterogeneity, as the SISO control performance rapidly degrades when the ignored neuron 2 is increasingly sensitive. Note that this effect is not symmetric, as discrepancies in sensitivity are substantially less problematic when neurons in the population are less sensitive to the light input as compared to the feedback neuron (relative gain*<*1). Taken together with previous multi-output modeling and estimation on experimental data, these simulations demonstrate that the techniques used first for SISO applications are readily applicable to multi-output problems and that such approaches could grant better control of population neural activity of interest.

## 4. Discussion

With the continued development of tools for precisely and selectively manipulating neuronal ensembles using multiple inputs ([13, 44]) and corresponding technologies for measuring large-scale neuronal activity ([1]), a framework for the integration of these technologies enables more intelligent interaction with neuronal circuits within and across brain regions (for review, see [14]). The state-space model structure is a natural choice for describing systems that involve a number of inputs and outputs (referred to as multi-input, multi-output or MIMO) ([45]). State-space models for system dynamics in combination with the framework of optimal control and estimation allows design and implementation of control loops to scale without cumbersome changes in methodology, as is evident in this study where the same model structure has been applied successfully in online estimation and control of activity in single-as well as multi-neuron systems.

In more modern control approaches, a model of the underlying dynamics to be controlled is used for both design and implementation of the control law. Across different pathways and circuits, numerous model types have been used to predict neural activity at fast time scales ([31, 46, 47]), some of which have been applied to ensembles of neurons ([28, 48–51]). For the purpose of control applications, we modeled optically-driven responses of neurons in the sensory thalamus using a state-space dynamical systems representation, where any higher order dynamics are captured in sets of coupled, first-order difference equations. In contrast to more widely used phenomenological models for neuronal responses to stimuli such as the linear receptive field ([52, 53]), the linear-nonlinear-Poisson (LNP) model ([42, 54]), and the generalized linear model (GLM) ([46, 48]), state-space models describe what could be large-scale recordings as arising from some potentially small number of latent ‘states’ that evolve dynamically in time as a function of themselves and covariates such as sensory or optical stimuli.

We found that linear state-space (*i.e*., GLDS) models could be used in the context of the control objective of maintaining a steady firing rate in the face of ongoing activity changes during wakefulness in these early sensory neurons. This was not a given, as many applications of state-space models to neuronal spiking data have used nonlinear dynamical systems, or at least linear dynamical systems with Poisson observations ([15, 18, 28, 30, 49, 55–58]). While the GLDS models used here clearly do not respect the statistics of the measured spiking data and while they can grossly mis-predict neuronal responses to optical stimulation (*e.g*., Figure 3(a)), they do capture the basic dynamics and the robustness of the parameter-adaptive Kalman filter in combination with feedback control allowed the use of this relatively simple modeling framework and unlocks a wealth of other linear design and analysis methods developed over years of study. Similarly, a recent study that used state-space feedback control in the context of hardware-simulated manipulation of electrocorticography (ECoG) made a practical concession to use a linear model that was more amenable to commonly-used feedback control techniques ([59]). In contrast, we previously used a linear-nonlinear-Poisson (LNP) spiking model in to design a classical proportional-integral (PI) controller ([7]). While that simple control strategy proved quite effective even for tracking patterns of rate modulation, the controller was designed numerically around a simulated spiking system, owing to the multiple nonlinearities that precluded the use of such design tools as LQR used in the present study. Simply put, there is a natural trade-off between the complexity of models and complexity of the control design and implementation itself, especially as the dimensionality of problems scale. That said, a nonlinear model will likely be needed in some control applications and it is possible the use of a PLDS model would have improved control performance even in this application but at the expense of complexity. In cases where a Poisson model is necessary and/or beneficial, there are previously-developed methods for estimating the underlying state of a PLDS (‘point process filter’, [56, 60]). One could leverage these nonlinear filtering techniques and design/implement feedback control in the log-linear state space as described here for linear systems.

Aside from the observed robustness to the linear approximation of nonlinear neuronal activity, we also found in the context of SISO control problems that the parameter-adaptive Kalman filtering and feedback control granted enough robustness to model-mismatch that the GLDS did not have to be fit during the tight time constraints of awake, head-fixed recording sessions. Instead, data from previous experiments were used for the controller design and online estimation (Figure 1(d)). Certainly, a large part of this success comes down to the fact that the control objective was relatively long-timescale firing rate regulation. In the context of trajectory tracking, we previously showed that, while closed-loop control grants some robustness to model mismatch, even things as simple as DC gain mis-estimation can lead to off-target activity when the target trajectory is time-varying ([7]). Indeed, in the present study we have observed a wide array of neuron sensitivity to light. It is also true that while the adaptive estimator was largely unbiased under SISO conditions, the parameter adaptation carried out here did not always eliminate bias in multi-output scenarios. Therefore, there are certainly scenarios like trajectory tracking and multi-output control in which the control objective would warrant better model fits, or at least adaptively re-estimating other parameters such as the input matrix (**B**) or output matrix (**C**) rather than attempting to capture all model mismatch with a linear disturbance as was implemented here. In general, this would call for nonlinear variants of the Kalman filter, such as the extended Kalman filter ([36, 60]).

In addition to the fact that the control method proved quite robust to model-mismatch in the sense of the statistics of measured data and the aforementioned long timescale biases in model predictions, we also found that we were able to effectively carry out the SISO control and estimation problems using a first order approximation of systems that appeared to be fourth or fifth order (Figure 2(b)). Since the control objective was to track a firing rate step command over relatively long timescales, this should be expected. After all, the dynamics of these systems tended to have died out after tens of milliseconds (Figure 2(b)). In applications where the objective is to entrain precisely-timed sequences of spiking activity rather than an overall firing rate (*e.g*., [18, 55, 61, 62]), a higher-order model would be merited and more emphasis would need to be placed on stimulus design.

The fact that the present study only tackles the problem of tracking a constant firing rate target begs the question of how applicable the approach is to the problem of tracking desired trajectories of neuronal activity. The controller was designed by solving an infinite horizon optimization problem (specifically, LQR) and was not explicitly designed for tracking target patterns of activity. That said, the methods laid out in here would be directly applicable to tracking problems where the desired pattern of activity is slow compared to the dynamics of the system being controlled. As mentioned above, the average light-to-spiking impulse response for thalamic units tended to die out over tens of milliseconds, indicating that the methods used here for model-based control may not be completely applicable to control of patterns at that time scale or faster. In such cases, a finite horizon optimization of feedback controller gains and a nominal control input using a technique like iterative LQR ([63]) may prove beneficial or necessary.

To this point, all references to the robustness of this control framework have pertained to activity of the putative single neuron which was used to adjust stimulation in real-time. Across recordings, the feedback neuron’s activity was maintained at the target firing rate with low error on average and, importantly, with low trial-to-trial variability. However, we found that the local population of neurons also excited by the optical stimulation did not exhibit this same lowered variability. It is worth noting that it is likely the case that pulsatile stimulation rather than the continuously-graded stimulation we used here would have had a less variable effect on the population. However, we showed that presenting 5 ms pulses of light synchronizes the population and would thus likely strongly impact downstream targets in a way that is unsuitable for many applications. Therefore, to reap the benefits of closed-loop control in neural circuits, feedback of population activity rather than a putative single neuron will likely be of great importance moving forward. Importantly, most commercially available electrophysiological data acquisition systems do not currently perform spike-sorting across dense multielectrode arrays like those used here; instead spike-sorting is carried out in a channel-by-channel fashion or is restricted to lower channel count tetrode configurations in general. Given the difficulty of online identification of individual neurons from raw electrophysiological recordings (*i.e*., spike-sorting) for dense multielectrode arrays, the thresholded multi-unit activity often utilized in brain-machine interface applications may prove an effective alternative measure of population activity ([64]). Alternatively, it is conceivable in the case of chronic implants to sort and track single units across experiments ([65, 66]).

Aside from providing multi-output feedback to the controller, the addition of multiple light sources (*e.g*., [44]) would afford some degree of population control spatially. The current preparation is highly underactuated in that there is a single light source being used to manipulate local activity, and there will in practice always be heterogeneities in responsiveness to light in space, whether it be due to varying distance from a common light source or differences in expression of opsins, *etc*. In addition to multiple spatially-distributed light sources, having the ability to simultaneously excite and inhibit neuronal activity using light of different wavelengths will also be key for robustness of optogenetic control moving forward. Note that in the present study, a single excitatory opsin (ChR2) was expressed in excitatory cells, meaning that the control is limited to pushing activity of those neurons toward higher firing rates. This effectively limits the control problem to one that maintains firing at an above-average desired level: here, 20 spikes/s which naturally occurs in this pathway. Conversely, if inhibitory opsins were expressed (or excitatory opsins were expressed in inhibitory interneurons), the control objective would be limited to maintaining or pulling down spontaneous levels of activity. Therefore, the ability to effectively push as well as pull back on neuronal activity would greatly expand the utility of this approach. Importantly, while not tested here, the state-space control and estimation methods developed in the present study should generalize to the control of more complex neural circuits in the future. However, it is likely the differing kinetics of excitatory and inhibitory opsins would necessitate higher order models.

Besides the utility in treatment of neurological disorders and diseases ([19, 20, 67, 68]), or in augmenting normal brain function, the precise, closed-loop control of neural circuits has the potential to significantly enhance our understanding of underlying mechanisms of basic brain function. After all, feedback control enabled the seminal work of Hodgkin and Huxley in uncovering the nature of the ionic currents that underlie the generation of a neuron’s action potential, for which they won the Nobel Prize in 1963 ([69]). The key to this experimental work was the use of a feedback controller to ‘clamp’ the trans-membrane voltage by injecting current to counter-act naturally occurring changes in ionic currents. This functional decoupling of constituent ionic and capacitive currents led to a quantitative description of the nonlinear dynamics of the action potential. Single-cell voltage clamp and dynamic clamp experiments ([70]) continue to be a powerful tool for scientific discovery, but continuously-graded feedback control of this sort has not been translated to the circuit-level, where the dynamics are complex and can adaptively change from moment to moment. Fundamentally different from lesioning or reversibly silencing brain regions, closed-loop optogenetic control has the opportunity to aid investigation of the mechanisms governing circuit level dynamics in a similar way voltage clamp did for the single neuron.

## 5. Conclusions

In this study, a state-space control and estimation framework has been developed and demonstrated to work well in the context of wakefulness, where there is spontaneous fluctuation in neuronal activity. Compared to Bolus *et al* ([7]), this updated approach is more naturally suited to the MIMO control problems that are important in studying complicated neural circuits. Notably, we were able to use a simple, linear approximation to this nonlinear system at least for long-timescale control objectives, such as maintaining an overall firing rate. The relative simplicity of these approaches achieved at the expense of modeling fidelity represents one of the chief strengths of the methodology, as linear control is well understood and is widely used. Moreover, while tested in the context of controlling spiking activity which is often statistically modeled as a point-process, the methods laid out here are immediately applicable to the control of continuous-valued neuronal signals of interest such as local field potential and voltage/calcium imaging. This demonstration of state-space models being used for single-and multi-output applications of optogenetic control opens the door to other established control strategies that use this modeling framework, such as model predictive control ([45]). As a whole, this work lays the foundation for future advances in manipulation and study of neuronal circuits using the integration of neuronal recordings and optogenetic stimulation.

## Acknowledgments

This work was supported by the NIH/NINDS Collaborative Research in Computational Neuroscience (CRCNS) / BRAIN Grant R01NS115327 (GBS and CJR) and NIH/NINDS BRAIN Grant R01NS104928 (GBS). MFB was supported by an NSF Graduate Research Fellowship Grant DGE1650044 and the Norman and Rosalyn Wells Fellowship. AAW was supported by the NIH/NIDA GT/Emory Computational Neuroscience Training Grant T90DA032466. CJR was additionally supported by NSF Grant CCF-1409422, and James S. McDonnell Foundation Grant 220020399.

## Notes

### Competing Interest Statement

The authors have declared no competing interest.

### Summary of Updates

This version has been updated to include additional information about the experimental and modeling process to increase clarity.

